# Keratin intermediate filaments mechanically position melanin pigments for genome photoprotection

**DOI:** 10.1101/2025.01.15.632531

**Authors:** Silvia Benito-Martínez, Laura Salavessa, Anne-Sophie Macé, Nathan Lardier, Vincent Fraisier, Julia Sirés-Campos, Riddhi Atul Jani, Maryse Romao, Charlène Gayrard, Marion Plessis, Ilse Hurbain, Cécile Nait-Meddour, Etienne Morel, Michele Boniotto, Jean-Baptiste Manneville, Françoise Bernerd, Christine Duval, Graça Raposo, Cédric Delevoye

## Abstract

Melanin pigments block genotoxic agents by positioning on the sun-exposed side of human skin keratinocytes’ nucleus. How this position is regulated and its role in genome photoprotection remains unknown. By developing a model of human keratinocytes internalizing extracellular melanin into pigment organelles, we show that keratin 5/14 intermediate filaments mechanically control the 3D perinuclear position of pigments, shielding DNA from photodamage. Imaging and microrheology in human disease-related model identify structural keratin cages surrounding pigment organelles to stiffen their microenvironment and maintain their 3D position. Optimum pigment spatialization is required for DNA photoprotection and rely on the interplay between intermediate filaments and microtubules bridged by plectin cytolinkers. Thus, the mechanically-driven proximity of pigment organelles to the nucleus is a key photoprotective parameter. Uncovering how human skin counteracts solar radiation by positioning the melanin microparasol next to the genome anticipates that dynamic spatialization of organelles is a physiological UV stress response.

**Short summary:** Melanin pigments shield DNA from photodamage by positioning atop nuclei in skin keratinocytes. We show keratin 5/14 intermediate filaments control this 3D spatialization, forming protective cages around pigments. This positioning, together with microtubule function, optimizes genome protection, revealing cytoskeletons and organelle dynamics as a UV stress response.

## INTRODUCTION

Human skin color and photoprotection rely on melanin pigments produced by melanocytes and then transferred to keratinocytes ^1^. This sequential two-cell type-dependent process occurs in the epidermis and consists, at least in part, of 1) melanin production in melanosomes, the lysosome-related organelles (LROs) of melanocytes ^2,3^; 2) transfer of melanin from melanocytes to keratinocytes ^4,5^; and 3) melanin packaging and perinuclear storage by keratinocytes ^1^. Although different modes of melanin transfer from melanocytes to keratinocytes have been documented ^1,6,7^, one process described in the human epidermis is the exocytosis of pigments by melanocytes followed by their phagocytosis by keratinocytes ^8–10^. Fusion of the limiting membrane of a melanosome with the plasma membrane of the melanocyte results in the extracellular release of its intraluminal melanin core, hereafter referred to as melanocore (MC). After internalization by keratinocytes, MCs reside in membrane-bound organelles ^8,9,11–13^ [MC-positive (MC^+^) organelles] whose cellular and molecular processes underlying their biology remain largely unknown ^1^.

In the human epidermis, keratinocytes sequester MCs in lysosome-like organelles carrying transmembrane proteins associated with organelles of the lysosomal lineage (i.e., LAMP1, CD63), while lacking tracers reflecting luminal acidity and degradative activity ^11,14^. The atypical molecular signature of the MC^+^ organelle has also been observed in a hybrid cell system using a mouse keratinocytic cell line (XB2) fed with pigment-enriched fractions isolated from the culture supernatant of a pigmented human melanoma cell line (MNT-1) ^15^. Although, to date, there is no characterized 2D *in vitro* model of human pigment-loaded keratinocyte ^5^, MC^+^ organelles in keratinocytes appear to be non-acidic, non-degradative lysosome-like compartments ^1^, proposed as a new member of the LROs family ^16^.

In situ, MC^+^ organelles are strategically positioned on the “sun-exposed” side of keratinocyte nuclei ^11,17–19^, in the form of a cap-like structure (a.k.a., microparasol) ^18,20,21^. The 3D perinuclear accumulation of melanin in keratinocytes is thus proposed to be critical for photoprotection by shielding their genetic material from the harmful effects of solar ultraviolet (UV) radiation. UV-induced DNA lesions include cyclobutane pyrimidine dimers (CPDs) and pyrimidine-pyrimidone (6-4) photoproducts (6-4PPs) ^22^, whose immuno-detection by fluorescence microscopy (FM) in human skin tissues of different phototypes is inversely correlated with the melanin content ^20,23–25^. Compared to lighter skin individuals, a lower incidence of photoinduced skin cancers is associated with darker skin individuals ^26,27^. In these individuals, proliferative basal keratinocytes contain a large amount of melanin pigments ^28^, and rarely show CPD-associated DNA lesions ^24,29^. Thus, and although skin photoprotection may in part rely on the amount of melanin pigments present in keratinocytes ^24^, we still ignore whether the intracellular position of the microparasol, and the cellular processes and molecular machineries governing the perinuclear distribution of MC^+^ organelles may be a key factor in genome photoprotection^1^.

The contribution of the cytoskeletons to the perinuclear distribution of pigments in keratinocytes remains poorly studied ^1,19^. While actin filament (F-Actin) dynamics contribute to pigment uptake by keratinocytes ^10,30^, microtubules (MTs) and the minus-end directed dynein motor were proposed to be involved in the long-range centripetal transport of pigments or bead particles ^18,19,31^. Genetic mutations responsible for haploinsufficiency of the keratin-5 ^32^ (*KRT5*)—a component of intermediate filaments (IFs)—are associated with the Dowling-Degos Disease (DDD: OMIM # 179850), a rare autosomal dominant genetic skin disorder characterized by hyperpigmented skin lesions in body folds, where a sparse distribution of melanin granules is observed in basal skin keratinocytes ^32–34^. In addition, loss-of-function mutations in other DDD-associated genes, such as *POFUT-1* or *POGLUT-1*, result in down-regulation of KRT5 expression ^35,36^. However, the contribution of IFs in the positioning of the MC^+^ organelle is not known.

IFs contribute not only to the cyto-protection against mechanical stresses ^37^, but also to organelle-related functions, such as their motion, immobilization, and thus positioning, through membrane binding or confinement ^38^. This is well illustrated for mitochondria, which can be surrounded by vimentin (VIM) or KRT5^+^ IFs that contribute to their position, morphologies, and metabolic functions ^38,39^. Cytoplasmic IFs form a highly dynamic and flexible filamentous network, extending from the nucleus to the cell periphery ^37^ and connecting to various intracellular structures, including F-actin and MTs cytoskeletons ^40,41^. Keratin components preferentially assemble in gene-regulated pairs differentially expressed according to the keratinocyte differentiation stage ^42^; e.g., KRT5 and its pairing partner KRT14 are specifically expressed in mitotically active basal keratinocytes ^43–47^. Yet, whether and how KRT5/14^+^ IFs contribute to the biology of MC^+^ organelles in keratinocytes have not been investigated.

Here, we address long identified but as yet unresolved questions of whether melanin organelles in keratinocytes are photoprotective and, if so, whether their 3D physical proximity to the nucleus is a key element. To this end, we developed and characterized a human cell-based system composed of MCs and primary human epidermal keratinocytes (HEKs). As in human skin, internalized MCs reside intact in perinuclear organelles with a non-acidic, non-degradative lysosome-like signature. The model demonstrates that MC^+^ organelles preferentially localized perinuclearly and atop the nucleus. To be retained in the perinuclear area, MC^+^ organelles specifically require intermediate filaments composed of KRT5 and KRT14 (KRT5/14^+^ IFs) that form cage-like structures, which partially encapsulate MC^+^ organelles and stiffen the mechanical properties of their microenvironment. Disruption of KRT5 expression softens the cytoplasm in a DDD model system, causing the dispersion of MC^+^ organelles, which is restored by KRT5 re-expression. Finally, optimal perinuclear 3D spatialization of MC^+^ organelles is necessary for DNA protection against UVB radiation through the cooperation between KRT5^+^ IFs and MTs bridged by the cytolinker plectin-1 (PLEC1) protein. Therefore, MC^+^ organelles are photoprotective structures, whose 3D positioning near and above the nucleus is a key parameter of their photoprotective activity. In conclusion, the mechanically induced pigment spatialization represents a physiological strategy developed by skin keratinocytes for cutaneous photoprotection.

## RESULTS

### Melanocores are the extracellular melanin contents of melanosomes

Prior to setting up an *in vitro* system to investigate the melanin logistics in human keratinocytes, we isolated and performed the first characterization of melanocores (MCs). The highly pigmented human melanoma MNT-1 cell line was chosen as a MCs-producing cell because it could be easily cultured in quantity and shared characteristics with normal human epidermal melanocytes (HEMs); e.g., expression levels of melanocyte-differentiated genes and protein products ^48,49^, and melanosome biogenesis ^50^. Based on established protocols ^15^, conditioned media from MNT-1 cells cultured for a week were collected, pooled, and centrifuged with concentrator-columns (see methods; **Extended Data Fig. 1a**, panels 1-2). The dark-brown MCs-enriched fractions (**Extended Data Fig. 1a**, panel 3; arrow) were pelleted (**Extended Data Fig. 1a**, panel 4) and their melanin concentration was measured by spectroscopy (**Extended Data Fig. 1b**) to show that MCs secreted by MNT-1 cells were consistently isolated in significant amounts.

The isolated MCs were first characterized at the ultrastructural level by transmission electron microscopy (TEM) (**Fig. 1a**). Like pigmented melanosomes ^51^, MCs were ellipsoidal and ∼0.5 μm in diameter (**Fig. 1a**, middle panels). Yet, unlike melanosomes, MCs lacked a limiting membrane (**Fig. 1a**, middle panels), as expected for secreted particles. MCs were also associated with small lipid vesicles of various electron densities decorating their surface (**Fig. 1a**, middle and right panels, arrowheads). These features were similar to MCs derived from highly pigmented primary HEMs (**Extended Data Fig. 1c**, arrowheads), indicating that MCs from MNT-1 cells are a physiologically relevant source of extracellular pigment.

**Figure 1.**
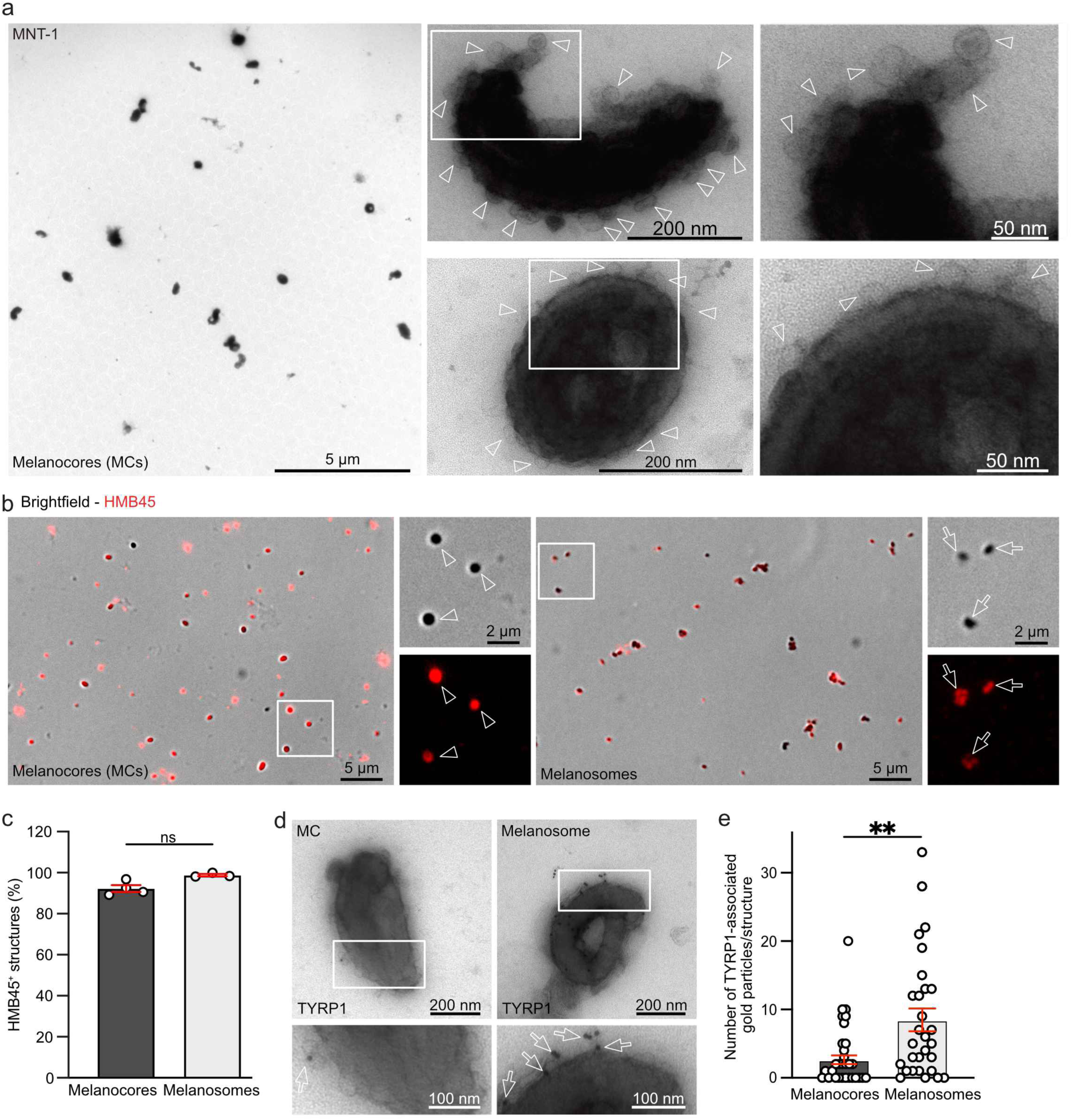
Melanocores are the secreted luminal melanin content of intracellular melanosomes. (**a**) EM micrographs of MCs isolated from MNT-1 cell culture medium showing pigmented particles ∼500 nm in size lacking limiting membranes and associated with lipid vesicles (arrowheads). (**b**) IFM of isolated MCs (left) or melanosomes (right) stained with HMB45 antibody (red, arrowheads and arrows, respectively) that appear as dark spots by brightfield illumination. (**c**) Average percentage of HMB45^+^-MCs or -melanosomes as in (b). (**d**) Immunogold labelling by EM of isolated MCs (left) or melanosomes (right) stained with TYRP1 antibody (arrows). (**e**) Average number of TYRP1-associated gold particles per pigmented structures. Data are the average of at least three independent experiments presented as the mean ± SEM. Two-tailed unpaired t test (ns, non-significant; *, *P* < 0.05).

Individual MCs were then quantitatively characterized by immuno-fluorescence microscopy (IFM) using a code that automatically detected dark pigments in the brightfield channel via a defined 1 µm region of interest centered around each MC, as well as their number per cell and their individual spatial coordinates (*x*, *y*, *z*) (see methods). Isolated MCs or pigmented melanosomes isolated by subcellular fractionation of MNT-1 cells homogenates were deposited on glass coverslips, chemically fixed and detected as black dots by brightfield microscopy (**Fig. 1b**). Isolated MCs and melanosomes were labeled with the HMB45 antibody (**Fig. 1b**, arrowheads and arrows, respectively; and **Fig. 1c**)—recognizing melanin-laden fibrils^5^. As previously observed by immuno-labeling by EM (IEM) of melanosomes ^5,50^, the darker MCs were the least labeled for HMB45 (due to the melanin deposits that covered the HMB45^+^ fibrils), and vice versa (**Extended Data Fig. 1d**). Consequently, the sub-population of optically-light HMB45^+^ MCs were not automatically detected in the brightfield channel (**Extended Data Fig. 1e**, arrowheads), and the code was modified (Code_1_; see methods) to detect all MCs populations, from the darkest to the lightest (**Extended Data Fig. 1f**, arrowheads). Finally, compared with isolated pigmented melanosomes, IEM analysis showed that MCs were poorly labeled for TYRP1 (**Fig. 1d**, arrows)—a melanin-synthesizing enzyme that is largely associated with the limiting membrane of melanosomes ^3^ and, to a lesser extent, with intraluminal vesicles of melanosomes in melanocytes ^50,52^. Whereas nearly half of the MCs analyzed were negative for TYRP1 (MCs: 43%; melanosomes: 17%), those that were positive decreased by 3-fold the number of gold particles associated with TYRP1 immunolabelling relative to melanosomes (**Fig. 1e**). Of these, almost a third of the TYRP1-associated gold particles decorated the small lipid vesicles associated with MCs (**Fig. 1d**, left panel, arrow). In conclusion, the data show that extracellular MCs of MNT-1 cells are a physiological source of pigment, collected in quantity and quality, that correspond to the secreted luminal content of pigmented melanosomes.

### Melanocores internalized by human keratinocytes are perinuclearly positioned in lysosome-like organelles

To develop a human pigmented keratinocyte model, MCs were synchronously deposited by centrifugation to a 2D-culture of primary human epidermal keratinocytes (HEKs). At early time points after deposition (0 and 10 min; **Fig. 2a** and **Extended Data Fig. 2a**, left and top panels, respectively), automatically detected MCs (arrowheads and dashed yellow circles) were predominantly peripheral and tightly associated with HEKs, and thus likely bound to their plasma membrane (PM; white dashed lines). At later time points (1 and 7 days; **Fig. 2a** and **Extended Data Fig. 2a**, right and bottom panels, respectively), MCs (arrowheads and dashed yellow circles) were intracellular and localized perinuclearly (as defined with DAPI-stained nuclei). The intracellular positioning of MCs over time was defined with Code_2_, which, combined with Code_1_, measured the approximate 2D-distance (d) along the x- and y-axes from each MC to the “nuclear envelope” (**Extended Data Fig. 2b**; see methods). MCs in HEKs positioned closer to the nuclear envelope over time (from 10 min to 7 days after deposition; **Fig. 2b**). Whereas MCs were located at an average distance of ∼14 µm at 10 min, this distance was significantly shortened by 5-fold at 8 h to approach the vicinity of the nucleus. One day after deposition, MCs reached an even shorter distance (2-fold decrease relative to 8h), maintained at 3 days, and even modestly reduced at 7 days.

**Figure 2.**
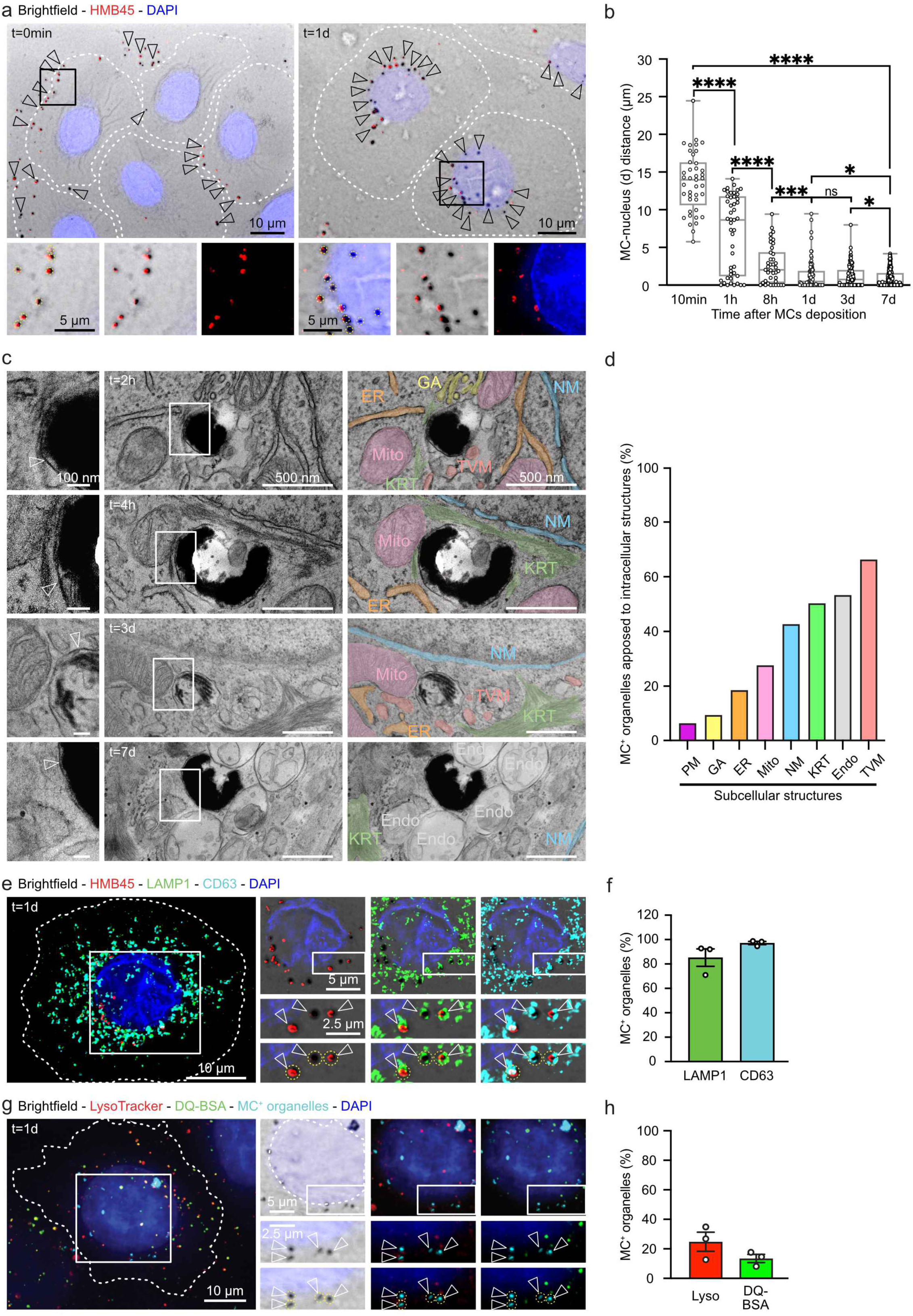
Internalized melanocores are perinuclearly positioned in lysosome-like organelles. (**a**) IFM of HEKs 0 min (left) or 1-day (right) after MCs (arrowheads) deposition and staining with HMB45 antibody (red) and DAPI (blue). (**b**) Quantification of the distance (d in µm) of each automatically detected MC as in (a) to their nucleus edge as a function of time after deposition. (**c**) Conventional EM micrographs of ultrathin sections of HEKs at different time after MC deposition. Insets (left) show MCs in single-membrane compartments (arrowheads) in proximity to various subcellular structures (right): keratin intermediate filaments (KRT; green), endoplasmic reticulum (ER; orange), mitochondria (Mito; pink), Golgi apparatus (GA; yellow), nuclear membrane (NM; blue), tubulo-vesicles (TVM; light red), endomembrane organelles (Endo; grey). (**d**) Quantification of the percentage of MC^+^ organelles in the vicinity of the subcellular structures shown in (c) in addition to plasma membrane (PM; purple). (**e**) SR-IFM of HEKs 1-day after MCs deposition and stained with HMB45 (red), anti-LAMP1 (green) and - CD63 (cyan) antibodies, and DAPI (blue). (**f**) Quantification of the mean percentage of MC^+^ organelles (as in e) positive for LAMP1 or CD63. (**g**) IFM of HEKs incubated with LysoTracker (red) and DQ-BSA (green) probes 1-day after MCs deposition (captured in brightfield and pseudo-colored in cyan) and stained with DAPI (blue). (**h**) Quantification of the mean percentage of MC^+^ organelles (as in g) positive for LysoTracker or DQ-BSA. Data are the average of at least three independent experiments presented as the mean ± SEM. Two-tailed unpaired t test (ns, non-significant; *, *P* < 0.05; ***, *P* < 0.001; ****, *P* < 0.0001). Automatically detected MCs (insets; arrowheads) are circled in yellow and cell outlines delineated by dashed lines.

We then used conventional EM to examine the cellular context associated with MCs. Consistent with optical imaging at early time points, extracellular MCs at 30 min (**Extended Data Fig. 2c**; arrowheads) were affixed to regions of the plasma membrane (left panel, arrows) that could form finger-like protrusions (right panel, arrow) likely contributing to MC internalization ^53^. Over time after deposition (2 h to 7 days), MCs were intracellular, surrounded by a limiting membrane (**Fig. 2c**, insets, arrowheads), and without ultrastructural alteration, showing together that internalized MCs reside intact in membrane-bound organelles ^14^, hereafter called MC^+^ organelles. As evidenced *in situ* ^11^, MC^+^ organelles showed proximity to various intracellular structures (**Fig. 2c**, middle and right panels)—the relative proportion of which was analyzed within a 500 nm radius distance from the center of each MC^+^ organelle (**Extended Data Fig. 2d**; and **Fig. 2d**)—such as plasma membrane (PM, magenta), Golgi apparatus (GA, yellow), endoplasmic reticulum (ER, orange), mitochondria (Mito, pink), nuclear membrane (NM, blue), intermediate filaments consisting of keratin bundles (KRT, green), endomembrane organelles (Endo, grey), or tubulo-vesicular membranes (TVM, light red).

In human epidermis or in 2D-cultured mouse keratinocyte cell lines, MC^+^ organelles were defined as non-acidic, non-degradative compartments carrying late endosomal/lysosomal membrane proteins (i.e., LAMP1 and CD63) ^11,15^. Using super-resolution IFM (SR-IFM) on HEKs one day after deposition (**Fig. 2e**), MC^+^ organelles stained with HMB45 antibody (red) were largely decorated by LAMP1 and CD63 (**Fig. 2e**, arrowheads, **Extended Data Fig. 2e**; and **Fig. 2f**). Next, live HEKs allowed to internalize MCs for 1 day were incubated with two fluorescent tracers probing either acidity (LysoTracker^TM^ Red) or degradative capacity (DQ^TM^ Green BSA) of cellular compartments, and then processed by FM (**Fig. 2g**). While both tracers were widely co-distributed in HEKs, as a reflect of their numerous acidic and degradative compartments, most perinuclear MC^+^ organelles (pseudocolored in cyan) were negative for both probes (**Fig. 2g**, arrowheads; and **Fig. 2h**).

In conclusion, primary human keratinocytes cultured *in vitro* internalize MCs, which are properly trafficked, positioned, and maintained for at least 1 week in non-acidic, non-degradative and perinuclear organelles bearing a lysosomal signature. Collectively, the human pigmented keratinocyte model recapitulates the intracellular localization, ultrastructural context, and molecular identity of MC^+^ organelles in human skin ^11^.

### Keratin-positive intermediate filaments encage melanocore organelles

Having set up a cellular model of pigmented keratinocytes with characteristics close to physiology, we first addressed how MC^+^ organelles were maintained in the perinuclear area, and then whether this position specifically contributed to keratinocyte photoprotection. Thus, we further examined the cytoskeletal elements in the immediate vicinity of MC^+^ organelles in keratinocytes. Ultrastructural study of pigmented HEKs identified that intermediate filaments (IFs), known to be composed of keratins ^45^, were frequently observed in the vicinity of MC^+^ organelles (**Fig. 2c-d**; green). Regardless of the skin color type, ultrastructural examination of HEKs fed with MCs (**Fig. 3a** and **Extended Data Fig. 3a**) or of keratinocytes in human pigmented skin epidermis (**Extended Data Fig. 3b**) revealed that keratin bundles often appeared to encapsulate MC^+^ organelles (arrows) in what we termed keratin cages (arrowheads). A main component of IFs in dividing basal keratinocytes is keratin-5 (KRT5) ^43,44^. IFM on HEKs 1 day after MCs deposition showed that KRT5 labeling formed a honeycomb-like pattern in the perinuclear area (**Fig. 3b**), likely reflecting numerous KRT5^+^ cages (yellow arrowheads); some of which were positive for MC^+^ organelles (white arrowheads). By conventional EM in HEKs, serial ultrathin sections of the same MC^+^ organelle showed that the KRTs^+^ cage did not fully surround the compartment in 3D (**Fig. 3a**, arrowheads in right panels). Consistently, 3D reconstruction from fluorescence imaging of a perinuclear portion of a HEK showed that MC^+^ organelles were apposed and partially surrounded by KRT5^+^ IFs (**Fig. 3c**, arrowheads and **Video 1**), together indicating that KRT^+^ cages are partially open structures around pigment organelles.

**Figure 3.**
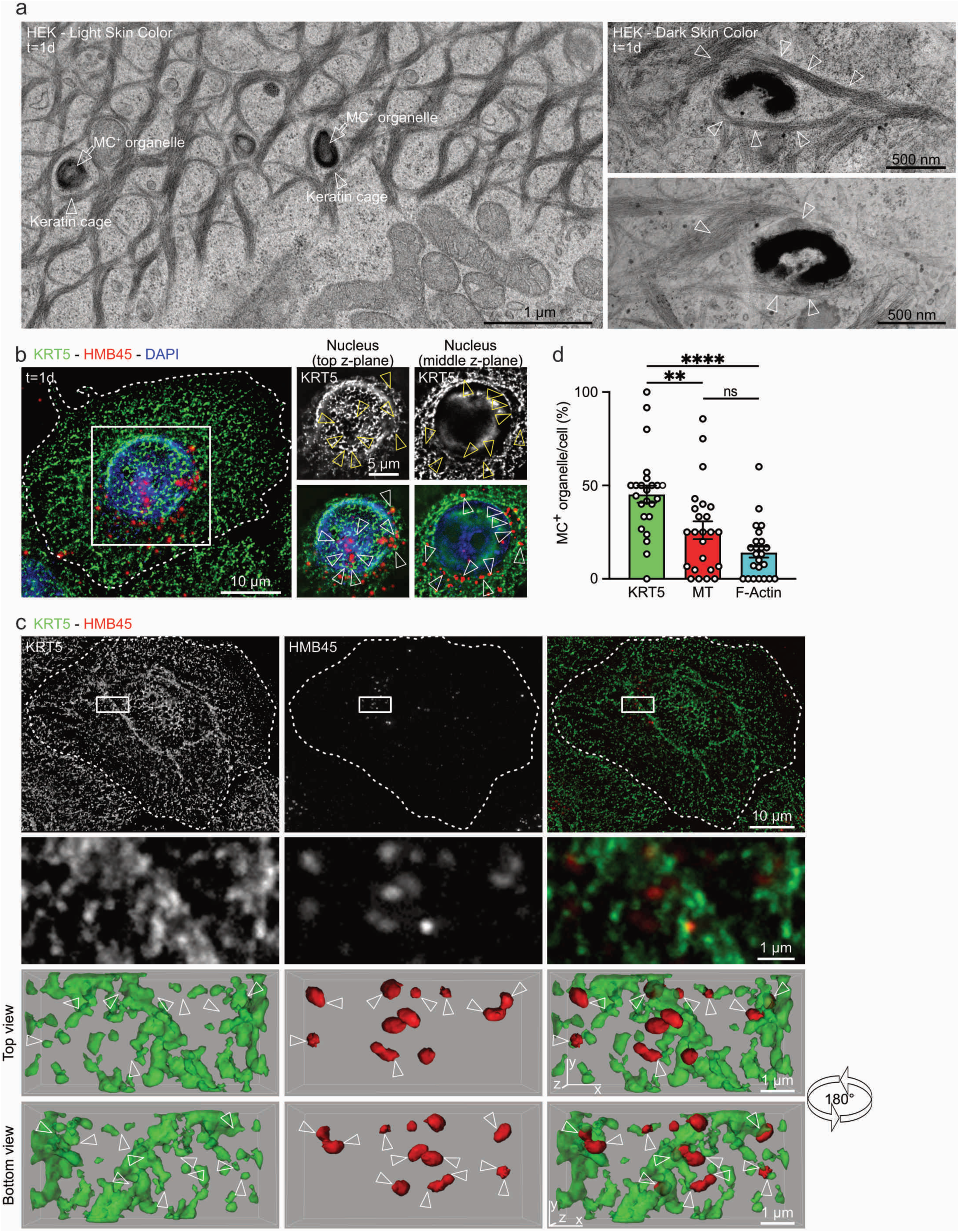
Melanocore organelles are surrounded by keratin 5 intermediate filaments in cage-like structures. (**a**) Conventional EM micrographs of ultrathin sections of HEKs 1-day after MCs deposition. Arrows point to MC^+^ organelles encircled by keratin^+^ intermediate filaments, named keratin cages (arrowheads). Insets (right) show the same MC^+^ organelle and its keratin cage (arrowheads) at two different z-planes. (**b**) IFM of HEKs 1-day after MCs deposition stained with KRT5 (green) and HMB45 (red) antibodies, and DAPI (blue). Insets (right) show MC^+^ organelles (arrowheads) at two different z-planes, at the top (left) or middle (right) of the nucleus. (**c**) IFM of HEKs 1 day after MCs deposition stained with anti-KRT5 (green) and HMB45 (red) antibodies and processed for 3D surface rendering of the boxed area to highlight the 3D organization of HMB45^+^ MC^+^ organelles in close proximity with KRT5^+^ IFs (arrowheads). Top and bottom views of the same area are presented (see also **Video 1**). (**d**) Quantification of the mean percentage of MC^+^ organelle per cell positive for KRT5, α-tubulin, or F-Actin staining (see also **Extended Data Fig. 3c-e**). Data are the average of three independent experiments presented as the mean ± SEM. Mann-Whitney two-tailed unpaired t test (ns, non-significant; **, *P* < 0.01; ****, *P* < 0.0001; ns, non-significant). The cell outline is delineated by a dashed line.

Next, we examined by IFM the overall subcellular distribution of different cytoskeletal elements in HEKs, namely KRT5^+^ IFs, MTs and F-Actin using antibodies against KRT5 or α-tubulin and fluorescence-conjugated phalloidin, respectively (**Fig. 3d**). Spatially, KRT5^+^ IFs and MTs were cytoskeletal elements more perinuclearly distributed than F-Actin (**Extended Data Fig. 3c**). Similar IFM on HEKs 1 day after MCs deposition (**Extended Data Fig. 3d-e**) showed that nearly half of MC^+^ compartments associated with KRT5^+^ IFs (similar to analysis by conventional EM, **Fig. 2d**), whereas this proportion significantly decreased 2-fold for MTs and F-Actin (**Fig. 3d**). Altogether, the data show that KRT5^+^ IFs in keratinocytes are cytoskeletal elements prominently associated with perinuclear MC^+^ organelles.

### Keratin 5/14 intermediate filaments are key for the perinuclear 3D-positioning of melanocore organelles

Because KRT5^+^ IFs cages partly encapsulate MC^+^ organelles, we investigated the role of KRT5 on their perinuclear position. First, we significantly decreased KRT5 protein expression level in HEKs using a pool of four distinct individual siRNAs (**Fig. 4a-c**). Then, after 1 day MCs deposition, IFM analysis of KTR5-depleted and DMSO-treated HEKs showed a 3-fold increase in the 2D-median distance (along x-/ y-axes) to the nucleus of MC^+^ organelles compared with control (**Fig. 4d** and **4g**). A pairing partner of KRT5 is KRT14 ^43–45,47^, and transiently reducing the expression of either of these KRTs in HEKs did not significantly perturb the expression of the respective partner (**Extended Data Fig. 4a-b**). Consistently, KRT14-depleted HEKs displayed a dispersion of the MC^+^ organelles along the x/y-axes of similar proportion as KRT5-depleted HEKs (**Extended Data Fig. 4c-e**). HEKs grown in culture express vimentin (VIM), an IFs protein co-existing with KRTs and contributing to their migration ^54^. As a control, HEKs were depleted of VIM, allowed to internalize MCs for 1 day, and similarly analyzed for MC^+^ organelles 2D-positioning (**Extended Data Fig. 4g-j**). No significant difference in the median distance to the nucleus of MC^+^ organelles or in the number of internalized MCs were observed between VIM- and CTRL-depleted HEKs (**Extended Data Fig. 4j-l**). Finally, and given that the average size of the nucleus was similar in all tested conditions (**Extended Data Fig. 4m**), the results show that KRT5/KRT14^+^ IFs specifically maintain MC^+^ organelles in the perinuclear area.

**Figure 4.**
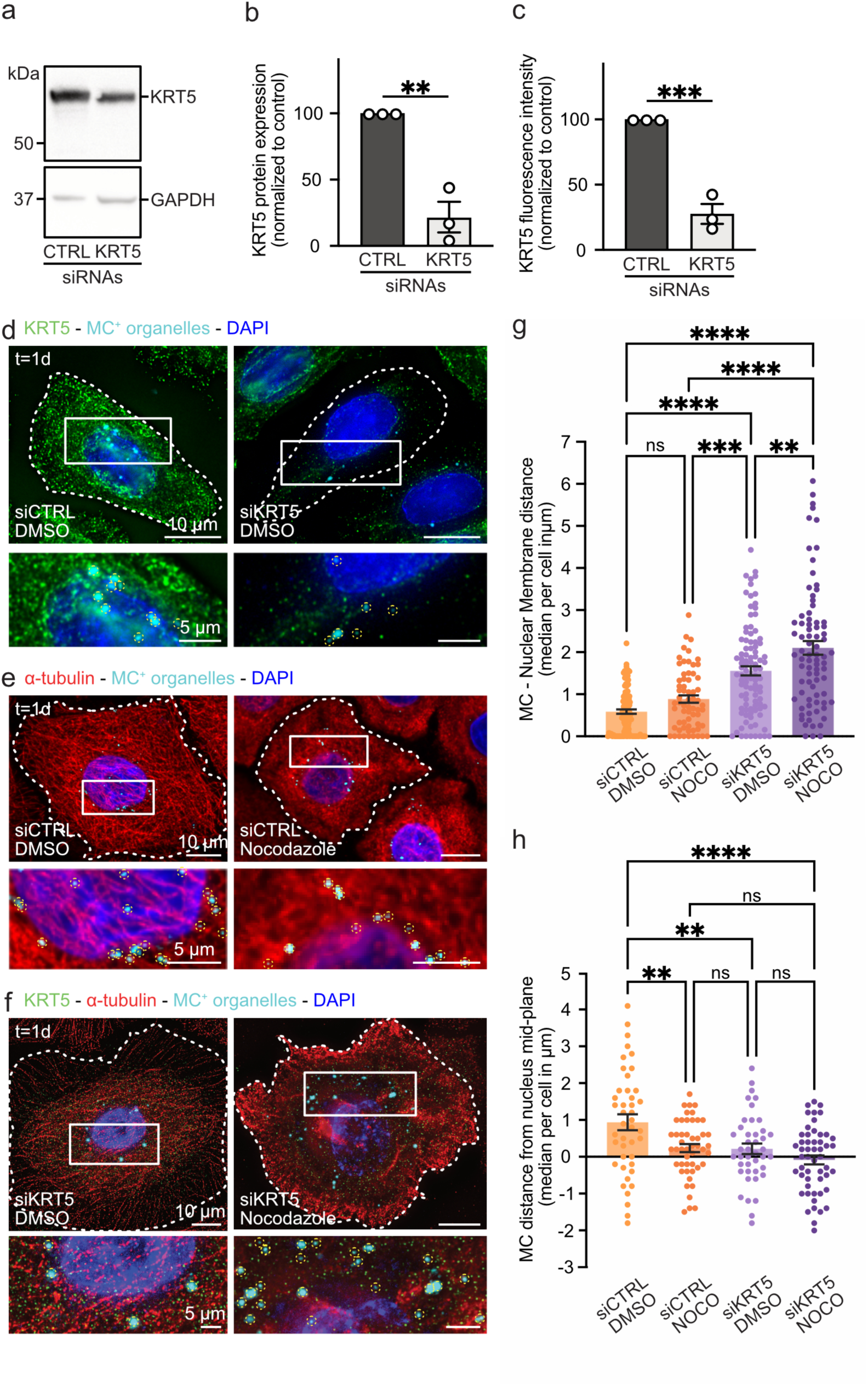
3D-position of melanocore organelles relies on KRT5^+^ intermediate filaments and microtubule network. (**a**) Western blot of HEK lysates treated with control (CTRL) or keratin 5 (KRT5) siRNAs and probed for KRT5 (top) or GAPDH (loading control, bottom). (**b**) Quantification of the average expression level of KRT5 in siCTRL- and siKRT5-treated HEKs normalized to GAPDH levels and control. (**c**) Quantification of the average fluorescence intensity of KRT5 in siCTRL- or siKRT5-treated DMSO-incubated HEKs used for analysis in figure 4d and normalized to control. (**d**) IFM of siCTRL-(left) or siKRT5-(right) treated HEKs incubated with DMSO and stained with KRT5 antibody (green) and DAPI (blue). (**e**) IFM of siCTRL-treated HEKs incubated with DMSO (left) or nocodazole (right) and stained with α-tubulin (red) antibody and DAPI (blue). (**f**) IFM of siKRT5-treated HEKs incubated with DMSO (left) or nocodazole (right) and stained with anti-KRT5 (green) and -α-tubulin (red) antibodies and DAPI (blue). (**g**) Quantification of the median distance of MC^+^ organelles to the nucleus edge in siCTRL- or siKRT5-treated HEKs incubated with DMSO or nocodazole expressed per cell. (**h**) Quantification of the median distance of MC^+^ organelles from nucleus mid-plane in siCTRL- or siKRT5-treated HEKs incubated with DMSO or nocodazole expressed per cell. (d-f) MCs were captured 1 day after deposition by brightfield microscopy and pseudo-colored in cyan. Automatically detected MCs are circled in yellow and cell outlines are delineated by dashed lines. Data are the average of at least three independent experiments presented as the mean or median ± SEM. Two-tailed unpaired t test and one-way ANOVA with Tukey post-hoc (ns, non-significant; *, *P* < 0.05; **, *P* < 0.01).

Since MTs likely support centripetal pigment transport in keratinocytes ^18,19,31^, we next addressed their contribution in maintaining the perinuclear position of MC^+^ organelles by incubating HEKs, after 1 day MCs deposition, with the MTs-depolymerizing drug nocodazole (NOCO). Compared with siCTRL-DMSO-treated cells, siCTRL-NOCO-treated HEKs showed a typical diffuse cytosolic α-tubulin staining (**Fig. 4e**) associated with a dispersion of the *trans*-Golgi network (TGN; labeled with TGN46 antibody) (**Extended Data Fig. 4n**). Surprisingly, NOCO treatment did not displace the 2D-position of MC^+^ organelles that remained as perinuclear as in DMSO-treated cells (**Fig. 4g**), showing that MTs do not significantly contribute to maintain the 2D-localization of MC^+^ organelles once they are in their perinuclear position. Then, we combined the siRNA-mediated depletion of KRT5 with the NOCO treatment and found a greater 2D-displacement of MC^+^ organelles from the nucleus, as compared to siKRT5-DMSO HEKs (**Fig. 4f-g**), demonstrating that MTs depolymerization amplifies the dispersion of MC^+^ organelles only in addition to KRT5 depletion.

In skin, MC^+^ organelles form a ‘pigmented microparasol’ above the nucleus, likely protecting the genome from UV rays. Thus, we used the human pigmented keratinocyte model to investigate the 3D-polarization of MC^+^ organelles. We measured the median z-axis position of MC^+^ organelles relative to the midplane of the nucleus and showed first that MC^+^ organelles were preferentially localized above the nuclear midplane in control HEKs. Second, MTs depolymerization or depletion of KRT5 or KRT14 reduced this vertical positioning by about 4- fold (**Fig. 4h** and **Extended Data Figure 4f**), with combined perturbations (siKRT5-NOCO condition) amplifying the effect, though not significantly compared with the control (**Fig. 4h**). Finally, depleting PLEC1, a cytolinker linking IFs to MTs ^40,41^, reduced by 2-fold the vertical positioning of MC^+^ organelles without affecting their 2D-distance to the nucleus (**Extended Data Fig 4o-t**). Together, these results show that the pigmented microparasol is formed *in vitro*, and that its polarization relies on the cooperation of two cytoskeletal systems. The KRT5/14^+^ intermediate filaments control the overall 3D-positioning of MC^+^ organelles, while microtubules, via PLEC1, contribute to refine their vertical positioning.

### Keratin 5 stiffens the microenvironment surrounding melanocore organelles

Next, we investigated whether KRT5^+^ IFs contribute to the perinuclear immobilization of MC^+^ organelles in keratinocytes, through local regulation of the mechanical properties of their cytoplasm microenvironment. As a pathophysiological relevant model, we used human keratinocytic cell lines (HaCaT) genetically engineered to carry the mutation in the head region of *KRT5* characterized in the Dowling Degos disease (*KRT5^DDD^*). While DDD is inherited as a dominant trait ^32^—such that patients are heterozygous for the mutation—here we generated homozygous HaCaT cells carrying the *KRT5^DDD^* mutation to study its consequences on pigment distribution in the pathophysiological context of DDD. As compared to parental cells (wild-type; WT), no KRT5 expression by IFM or immunoblotting was detected in HaCaT KRT5^DDD^ cells (**Fig. 5a**, bottom panel; and **Extended Data Fig. 5a**), whereas KRT14 was similarly expressed in both cells (**Extended Data Fig. 5a**). Therefore, HaCaT KRT5^DDD^ cells could be considered as KRT5 KO cells.

**Figure 5.**
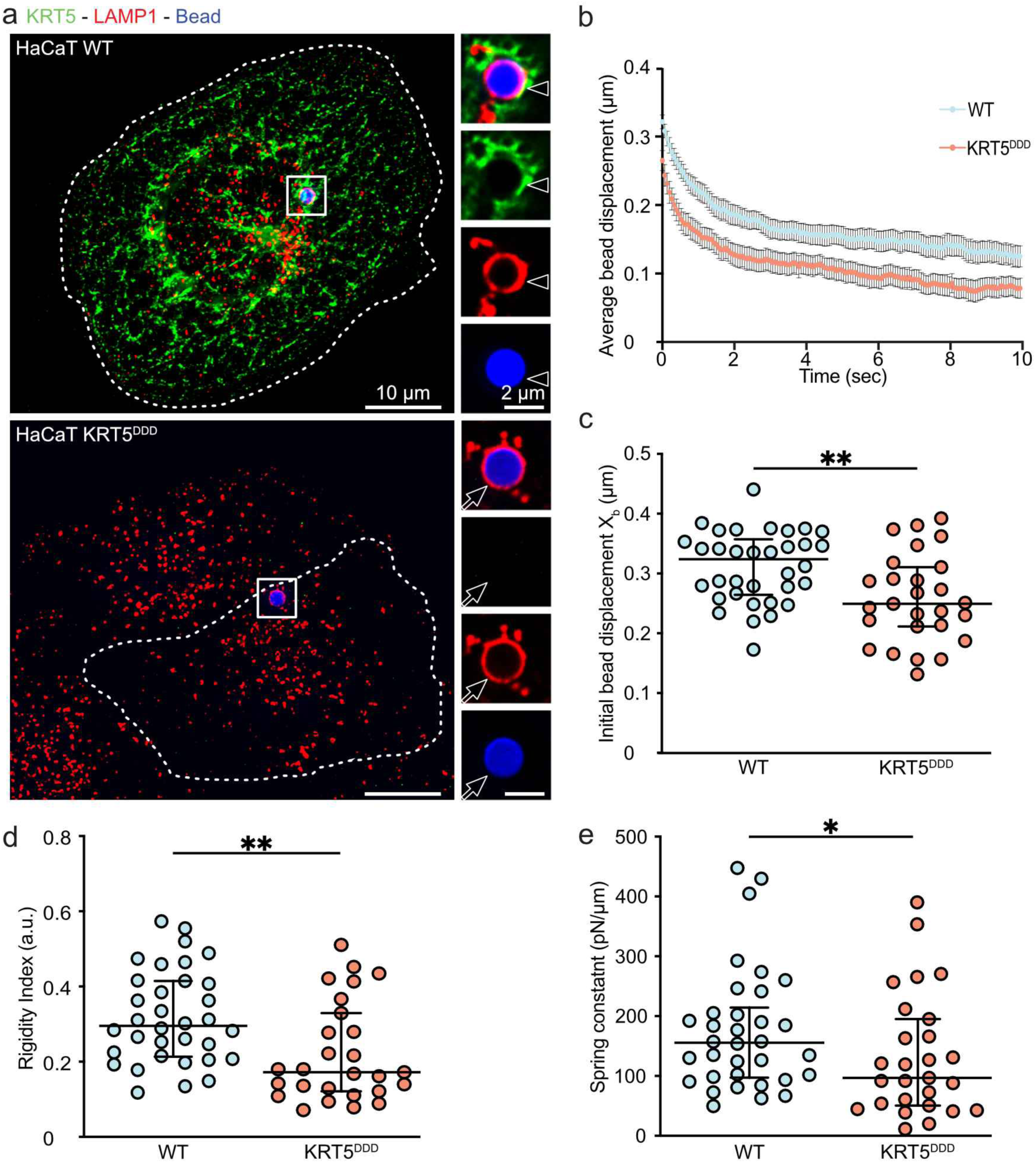
The Dowling-Degos disease keratin 5 mutation decreases the stiffness of the cytosolic microenvironment. (**a**) IFM of HaCaT cell wild-type (WT, top) or carrying the DDD-associated *KRT5* mutation (KRT5^DDD^, bottom) having internalized 2-μm beads (blue) and stained with KRT5 (green) and LAMP1 (red) antibodies. Arrowheads (top, insets) point to KRT5 staining surrounding internalized bead in a WT cell, and arrows (bottom, insets) point to the absence of KRT5 signal in a KRT5^DDD^ cell. Cell outlines are delineated by dashed lines. (**b**) Quantification of the average bead displacement (in µm) in WT (blue) or KRT5^DDD^ (red) HaCaT cells as a function of time (in sec) showing viscoelastic relaxation of the bead towards the trap center following a 0.5 μm step displacement of the microscope stage. (**c**-**d**) Quantification of the relaxation curves (shown in b) using a phenomenological model approach giving the initial bead displacement (c; or bead-step amplitude, X_b_ in µm) and the rigidity index (d) in WT (blue) or KRT5^DDD^ (red) HaCaT cells. (**e**) Quantification of the relaxation curves (shown in b) using the Standard Linear Liquid (SLL) viscoelastic model and analysis of the elasticity (spring constant in pN/µm) of the cytosolic microenvironment of the bead in WT (blue) or KRT5^DDD^ (red) HaCaT cells. Data are the average of at least three independent experiments presented as the mean ± SD. Mann-Whitney two-tailed unpaired t test (*, *P* < 0.05; **, *P* < 0.01).

First, HaCaT KRT5^DDD^ cells were used to directly assess the contribution of KRT5 in positioning MC^+^ organelles, by re-expressing KRT5 fused to mCherry (KRT5-mCh) in cells that internalized MCs for 1 day, followed by nocodazole treatment to eliminate the contribution of MTs. Expression of KRT5-mCh partially restored the perinuclear MC^+^ organelles distribution (**Extended Data Fig. 5b**), demonstrating that KRT5 is sufficient for this positioning. Second, live cell imaging of HaCaT KRT5^DDD^ cells—incubated with SiR-Tubulin to label microtubules— showed that, compared to WT, tracked MC^+^ organelles were more mobile, explored larger areas and moved faster, with some movements aligned with microtubules (**Extended Data Fig. 5c** and **Videos 2-3**), indicating partial microtubule-based movement of MC^+^ organelles in the absence of KRT5 expression. Third, we tested the contribution of KRT5 to the mechanical properties of the cytoplasm microenvironment by using KRT5^DDD^ HaCaT cells, in contrast to siKRT5-treated HEKs that only showed a partial reduction in KRT5 expression (**Fig. 4a-c, Extended Data Fig. 4a-b**).

We probed the cytoplasm viscoelastic properties in live cells by using optical tweezers-based active microrheology. Viscoelastic relaxation following a 0.5 µm step displacement of a 2 µm-diameter bead, internalized in either WT or KRT5^DDD^ HaCaT cells, and trapped by optical tweezers allowed to measure the deformability of the cytoplasm upon loss of KRT5 expression (see methods; and **Extended Data Fig. 5d**). The 2 µm beads were similarly internalized by both HaCaT cell types where they localized in LAMP1/CD63^++^ compartments (**Fig. 5a** and **Extended Data Fig. 5e**), as previously described for retinal pigment epithelial cells ^55^. In WT HaCaT cells, the internalized beads were embedded in a perinuclear KRT5 network (**Fig. 5a**, arrowheads; and **Extended Data Fig. 5f**), which was absent in KRT5^DDD^ cells (**Fig. 5a**, arrows). Therefore, internalized beads distribute in perinuclear lysosomal-like compartments surrounded by KRT5^+^ IFs, which represent a proxy for the subcellular distribution and environment of MC^+^ organelles.

Rheological measurements showed that the beads relaxation towards the optical trap center was faster in KRT5^DDD^ cells than in WT cells (**Fig. 5b**), as shown by the greater negative slope in KRT5^DDD^ cells at t=0 sec (**Fig. 5b**) that indicates a softer perinuclear cytoplasm. Analysis of the relaxation curves by a phenomenological approach (**Extended Data Fig. 5d**) showed that the initial displacement of the beads following the 0.5 µm step (**Fig. 5c**) and the rigidity index (**Fig. 5d**, see methods) were significantly reduced in KRT5^DDD^ cells, confirming the greater deformability of their cytoplasm compared with that of WT cells. Finally, results were refined by a viscoelastic analysis using the Standard Linear Liquid (SLL) model (**Extended Data Fig. 5d**), which demonstrated that the elasticity (spring constant) of the cytoplasm was significantly decreased in KRT5^DDD^ cells (**Fig. 5e**), while the viscosity was not affected (**Extended Data Fig. 5g**). Together, the data show that KRT5 contribute to the elasticity of the perinuclear cytoplasm in keratinocytic cells, suggesting that KRT5^+^ IFs immobilize MC^+^ organelles perinuclearly by increasing the local stiffness of its microenvironment.

### Genome photoprotection relies on the perinuclear 3D-position of melanocore organelles

In the human epidermis, the perinuclear accumulation of pigments in basal keratinocytes, a.k.a., the microparasol, presumably protects their genetic material from the harmful effects of solar UVB radiation ^23,56,57^. *In vitro*, the incorporation by HEKs of melanins derived from detergent-treated melanosomes provided a certain level of DNA photoprotection ^58^. However, it is still not known whether genome photoprotection depends on the spatialization of the microparasol, and therefore on the perinuclear 3D-localization of MC^+^ organelles. To test this hypothesis, we first investigated whether the newly established human keratinocyte-based system was suitable for monitoring UVB-dependent DNA photodamage. HEKs (not incubated with MCs) or HEKs allowed to internalize MCs for 1 day were exposed to UVB doses (0.5 or 1 J/cm^2^), and immediately processed by IFM using an antibody against cyclobutane pyrimidine dimers (CPDs), a UVB-dependent DNA photolesion. HEKs devoid of internalized MCs and exposed to both UVB doses showed robust nuclear staining for CPDs (**Fig. 6a**, top panels). In contrast, the mean fluorescence intensity of the nuclear CPD staining was significantly reduced in UVB-exposed HEKs that had perinuclear MCs (**Fig. 6a**, bottom panels; and **Fig. 6b**). For both UVB doses, the mean nuclear CPD fluorescence intensity was inversely correlated with that of the HMB45 signal in the nucleus region of interest (**Extended Data Fig. 6a**), showing that the more MCs are present in the nuclear area, the fewer UVB-dependent DNA photolesions. Collectively, the data show that perinuclear MC^+^ organelles in HEKs are intracellular compartments with DNA photoprotective function.

**Figure 6.**
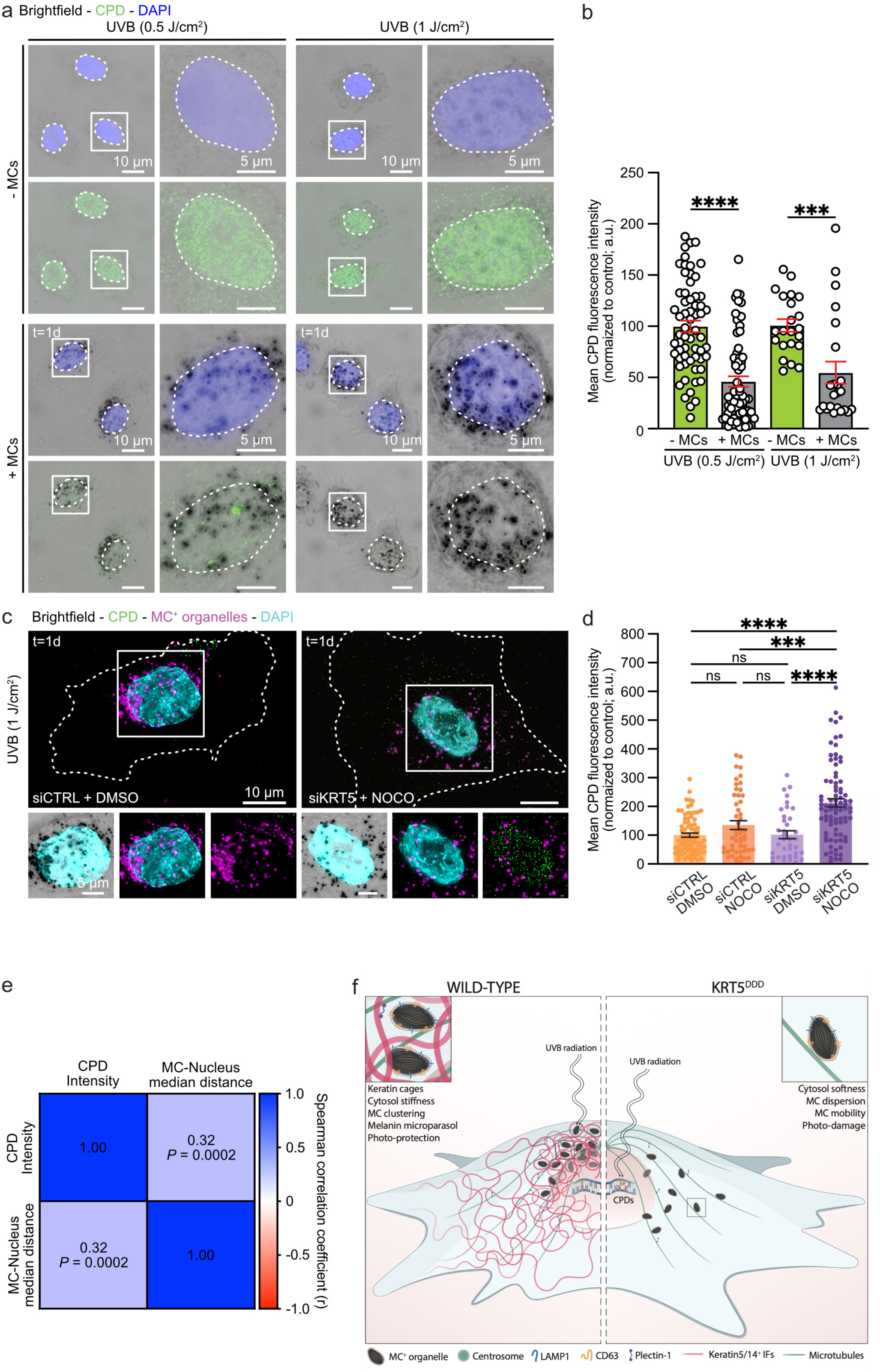
DNA photoprotection relies on melanocores organelles and their perinuclear 3D-position. (**a**) IFM of HEKs with (bottom) or without (top) 1-day deposition of MCs exposed to a UVB dose of 0.5 J/cm^2^ (left) or 1 J/cm^2^ (right) before staining with CPD antibody (green) and DAPI (blue). MCs were captured by brightfield microscopy and nucleus outlines delineated by dashed lines. (**b**) Quantification of the mean nuclear CPD fluorescence intensity in cells treated as in (a) and normalized to respective controls. (**c**) IFM 1-day after MCs deposition of DMSO siCTRL-(left) or nocodazole siKRT5-(right) treated HEKs exposed with 1 J/cm^2^ UVB dose before staining with CPD (green) and DAPI (cyan). MCs captured by brightfield microscopy were pseudo-colored in magenta. Cell outlines are delineated by dashed lines. (**d**) Quantification of the mean nuclear CPD fluorescence intensity in cells treated as in (c) and normalized to control. (**e**) Matrix of correlation between the mean nuclear CPD fluorescence intensity and the mean MC organelle-nucleus median distance of siCTRL-DMSO and siKRT5-NOCO-treated HEKs and showing the non-parametric Spearman correlation coefficient (r). (**f**) Working model on the role of KRT5/14^+^ IFs (pink) in the perinuclear positioning of MC^+^ organelles and UVB-dependent DNA photoprotection in skin keratinocytes. (**Left**) Under normal conditions, extracellular MCs are internalized and stored in non-acidic and non-degradative MC^+^ organelles carrying CD63 and LAMP1 (yellow and blue, respectively) located in the perinuclear area atop the nucleus to form a melanin microparasol shielding DNA from UVB-dependent damages. There, MC^+^ organelles are surrounded by KRT5/14^+^ IFs in cage-like structures that mechanically maintain their 3D-proximity to the nucleus by rigidifying the cytosolic microenvironment, while microtubules (green) aid in their vertical positioning via plectin 1-dependent bridging to IFs (purple). (**Right**) Loss of KRT5 expression or presence of the Dowling-Degos disease (DDD)-associated *KRT5* mutation leads to the dispersal of MC^+^ organelles due to cytosol softening and increased microtubule-dependent motility. Disrupting the microparasol by altering KRT5^+^ IFs and microtubules enhances the 3D-dispersion of MC^+^ organelles, resulting in more UVB-induced DNA photolesions (CPDs) showing suboptimal photoprotection of genetic material from UVB solar radiation. Data are the average of at least three independent experiments presented as the mean ± SEM. One-way ANOVA statistical analysis with Tukey post-hoc (ns, non-significant; **, *P* < 0.01; ***, *P* < 0.001; ****, *P* < 0.0001).

Finally, we tested if the perinuclear 3D-spatialization of MC^+^ organelles was crucial for their photoprotective activity. We treated HEKs, allowed to internalize MCs for 1 day, with KRT5 siRNAs, or NOCO, or both, and exposed them to 1 J/cm^2^ UVB, before CPD immunolabeling (**Fig. 6c**; **Extended Data Fig. 6b**). In all conditions, HEKs were fed with a similar number of MCs (**Extended Data Fig. 6c**) to now eliminate the contribution of the amount of melanin pigments per cell in the photoprotective assay (**Extended Data Fig. 6d**). Then, the mean nuclear CPD fluorescence intensity significantly doubled in siKRT5 NOCO-treated HEKs compared with other conditions (**Fig. 6d**), highlighting that genome photoprotection relies *in vitro* on both KRT5/14^+^ IFs and an intact MTs network to ensure a 3D-spatialized and functional microparasol. Consistently, the positive correlation between the MC-nucleus median distance and the nuclear CDP fluorescence intensity (**Fig. 6e**) showed that the greater the distance between MCs and the nucleus, the higher CPD fluorescence intensity in the nucleus. Together, MC^+^ organelles in human keratinocytes are intracellular compartments whose perinuclear 3D-positioning and short distance to the nucleus are key for the UVB-dependent DNA photoprotection (**Fig. 6f**).

## DISCUSSION

Melanin pigments provide the pigmentation of human skin and natural photoprotection. The cellular and molecular processes underlying melanocyte pigmentation have been extensively studied ^3^, but those governing pigment fate in human keratinocytes remain comparatively unexplored ^1^. To bridge this gap, we report here an *in vitro* human pigmented keratinocyte system consisting of primary HEKs and internalized MCs isolated from human pigmented melanoma cells. The model recapitulates the features of pigmented keratinocytes described in human skin ^8,11^; i.e., extracellular MCs (**Fig. 1**) are internalized by HEKs and reside intact in lysosome-like organelles (MC^+^ organelles), positive for LAMP1 and CD63, but neither acidic nor degradative, and which form a pigmented microparasol by being positioned atop nuclei (**Fig. 2**, **Fig. 4**). Perinuclear 3D-polarization of MC^+^ organelles is required for DNA photoprotection against UVB (**Fig. 6**) and relies on KRT5/14^+^ IFs and microtubules bridged by plectin 1 (**Fig. 4**, **Fig. 6**). KRT5^+^ IFs form cage-like structures (**Fig. 3**) that can immobilize pigments through local confinement and stiffening of their microenvironment, which is affected in a DDD model system (**Fig. 5**). Collectively, the new model uncovers that the pivotal perinuclear 3D-positioning of pigment organelles is a physiological photoprotective strategy of keratinocytes, orchestrated by the interplay of two cytoskeletal elements, the keratin intermediate filaments and the microtubules (**Fig. 6f**).

Melanocores secreted by melanocytes, which correspond to the luminal content released from intracellular melanosomes, have not been extensively characterized ^15^. Here, MCs released from highly pigmented human melanoma cells are similar to those of primary human pigmented melanocytes, i.e., MCs are non-membrane-bound ellipsoidal melanin particles of half µm-size decorated with membranous vesicles that carry a cohort of the melanin-synthesizing enzyme TYRP1 (**Fig. 1**, **Extended Data Fig. 1**). These vesicles likely reflect the intracellular origin of MCs in melanocytes, given the association of MCs with HMB45^+^ melanized fibrils, whose formation depends on intraluminal vesicles formed in melanosomes ^59,60^. Indeed, TYRP1^+^ intraluminal vesicles surrounding the intracellular melanin cores are present in melanosomes ^50,52^, but it cannot be ruled out that similar vesicles secreted by pigment cells and originating either from the endosomal system (exosomes) or the plasma membrane (ectosomes)—and collectively referred to as extracellular vesicles (EVs) ^61^—may also associate with MCs once present in the extracellular space. Finally, although the role of MC-associated vesicles is not known, they could contribute to pigment uptake and/or signaling in keratinocytes such as during EVs-mediated intercellular communication ^61^.

In human skin and in mouse keratinocytic cells, the internalization of MCs could occur by phagocytosis ^8,10^. As MC^+^ organelles persist for weeks in human skin, they could consist of phagosome-like compartments, which would not be canonically targeted for degradation ^14^. In contrast to the luminal content of phagosomes that is degraded within a few hours, following fusion with lysosomes and the formation of acidic and degradative phagolysosomes ^62^, MC^+^ organelles are long-lived structures that remain *in vitro* intact in HEKs for at least a week (**Fig. 2**). Although positive for LAMP1 and CD63 late-endosomal/lysosomal components, perinuclear MC^+^ compartments are neither acidic nor degradative (**Fig. 2**) such as in human skin ^11^. These components could be acquired by fusion of lysosomes with maturing stages of MC^+^ organelles prior to acquisition of their perinuclear position ^14^. Finally, organelles characterized by a lysosomal signature without degradative capacity and/or low pH are reminiscent of members of the LROs family, to which MC^+^ organelles may belong ^16^.

In human skin, and particularly in basal epidermal keratinocytes, pigments accumulate above the nuclei of keratinocytes ^11,17,63^, the so-called nuclear microparasol. Our microscopy data on 2D-cultured HEKs show that MC^+^ organelles are preferentially positioned in the perinuclear area atop the nucleus (**Fig. 2**, **Fig. 4**; and **Extended Data Fig. 3-4**). As observed in human skin ^11^, they are affixed not only to the nuclear envelope and other organelles, such as the endoplasmic reticulum or mitochondria (**Fig. 2**), but also to IFs (**Fig. 3**, **Extended Data Fig. 3**)^64^. We observe that KRT5/14^+^ IFs form a perinuclear honeycomb-like network, where MC^+^ organelles lodge in alveoli. This reticulated network is not limited to KRT5/14^+^ IFs, as vimentin (VIM) can form filamentous cage-like structures surrounding organelles, such as lipid droplets, the nucleus, or pigment organelles in amphibian pigment cells ^65–67^. Although the organelle-related functions of IFs cages are not clearly understood, they would confine enclosed compartments and isolate them from the cytosolic environment, likely influencing their position, shape, motility, membrane dynamics and contacts, and/or functions ^37,38^. For instance, the KRT6/16 pair contributes to maintain mitochondria morphology, dynamics, and function in mouse keratinocytes ^68^. Here, KRT5 or its partner KRT14, but not VIM, is required for the 3D-perinuclear maintenance and immobilization of MC^+^ organelles (**Fig. 4**, **Extended Data Fig. 4-5**), showing that a specific subset of IFs is required for MC^+^ organelles positioning and dynamics. However, it remains to be investigated if the KRT5/14^+^ IFs cages could be dynamic structures impacting the capture and/or the ability of MC^+^ organelles to contact intracellular organelles or membranes such as the nuclear envelope ^38,69^.

IFs are flexible and robust cytoskeletal components with peculiar elastic properties, which help cells cope with mechanical stress ^37^. For example, keratins contribute to intracellular mechanics, notably by stiffening the cytoplasm in the perinuclear area ^70^. The introduction of the *KRT5* mutation characterized in DDD specifically resulted in a near complete loss of KRT5 expression and caused reduced cytoplasmic elasticity (**Fig. 5**, **Extended Data Fig. 5**). These results are consistent with the superelastic characteristics of IFs as proposed for those composed of VIM or keratins ^71,72^, and the decrease in cell stiffness in absence of keratin networks ^73,74^. We defined the local contribution of KRT5 to the biophysical parameters of the cytosol by using optical tweezers and 2 µm beads. Although larger than MCs, beads are internalized in HaCaT cells and similarly packaged in LAMP1/CD63^++^ perinuclear organelles surrounded by KRT5^+^ IFs (**Fig. 5**, **Extended Data Fig. 5**). Because MC^+^ organelles were difficult to capture and keep intact in the optical trap, likely due to their smaller size and the natural optical absorption properties of melanin ^5^, the 2 µm beads represent a proxy for MC^+^ organelles. Considering the specific dispersion of MC^+^ organelles in HEKs with reduced KRT5 or KRT14 expression and the restoration of their perinuclear distribution in KRT5^DDD^ HaCaT cells expressing KRT5-mCh (**Fig. 4**, **Extended Data Fig. 4-5**), the biophysical results suggest that KRT5/14^+^ IFs increase the local stiffness of the MC^+^ organelles microenvironment, likely via IFs cage-like structures, to cause their maintenance and their immobilization near the nucleus.

Beyond their biophysical contribution, KRT5/14^+^ IFs could modulate the intracellular distribution of MC^+^ organelles by binding to their membrane as proposed for mitochondria ^38^, and/or by physically connecting them to the nuclear envelope ^69^. In addition, KRTs^+^ IFs can functionally co-exist with other cytoskeletons. This is the case for VIM which interacts with KRT14 to regulate the migratory phenotype of keratinocytes ^54^. However and given that VIM did not contribute in HEKs to the intracellular distribution of MC^+^ organelles (**Extended Data Fig. 4**), KRT5/14 are very likely the major components of IFs required for the 3D-maintenance of the pigmented microparasol. The IFs also establish crosstalk with the actin and microtubule networks ^37^, via cytolinkers like plectins ^40,41^. While we have not examined the contribution of actin in the positioning of MC^+^ organelles, our results (**Fig. 4**, **Fig. 6**) indicate that KRT5/14^+^ IFs do not contribute to the internalization of MCs unlike F-Actin ^10^. Importantly, MTs depolymerization or reduced PLEC1 expression only affected the vertical positioning of MC^+^ organelles by shifting them downward relative to the nucleus midplane, whereas combined disruption of MTs and KRT5^+^ IFs amplified the overall 3D-dispersion compared with altering KRT5 expression alone (**Fig. 4**, **Extended Data Fig. 4**). This proposes that KRT5/14^+^ IFs are crucial for the 3D-spatial organization of the microparasol, while MTs, via PLEC1-dependent bridging to IFs, fine-tune its height relative to the nucleus. Together, these findings support that these two cytoskeletons coordinate their function yet operate independently. Consistently, re-expression of KRT5 in MTs-disrupted KRT5^DDD^ HaCaT cells restored the perinuclear localization of MC^+^ organelles (**Extended Data Fig. 5**). Interestingly, MTs and centrosome-related machinery may help orient the melanin nuclear cap in human skin explants ^19^, suggesting that 2D- and 3D-organized keratinocytes exploit MTs, IFs and the way they interact to polarize perinuclear MC^+^ organelles above the nucleus.

Collectively, the results lead us to propose a model (**Fig. 6f**) in which MCs are internalized and packaged in MC^+^ organelles that are then transported retrogradely along MTs to the perinuclear area, where they are immobilized and isolated from their microenvironment due to the physical constrains provided by KRT5/14^+^ IFs and their cage-like structures. In this context, KRT5/14 play a crucial role in forming and maintaining the pigmented microparasol by regulating its 3D spatial arrangement within the cellular framework, while MTs together with plectin cytolinkers fine-tune its polarization above the nucleus. Consistently, previous *in vitro* models exploiting keratinocytic cells having internalized melanosomes or beads have shown that both particles would be transported centripetally along MTs using the dynein motor ^18,31^. The 3D-positioning of MC^+^ organelles by KRT5/14^+^ IFs and MTs therefore contributes to strategically place the melanin microparasol as close as possible to the UV-exposed side of the nuclear genome, maximizing its optical absorption property for optimal photoprotection (**Fig. 6f**). Conversely, a failure to position and immobilize MC^+^ organelles in the 3D-perinuclear area via the perturbation of the KRT5/14^+^ IFs cages and MTs dynamics or interaction with IFs can lead to suboptimal photoprotection, due to their increased distance from the genetic material (**Fig. 6f**).

Melanin nuclear-caps are thought to protect the keratinocyte genome against UV-induced DNA damage ^1^, but whether the perinuclear location is a determining photoprotective factor is not known. As defined *in vitro* (**Extended Data Fig. 6**), studies on human skin sections show that the amount of melanin in keratinocytes is inversely correlated with DNA photoproducts; i.e., they are higher in lightly pigmented skin and lower in highly pigmented skin ^20,24^. In addition, *in vitro* approaches exploiting HEKs fed with melanin extracted from detergent-treated melanosomes have shown that such incorporated melanin can provide DNA photoprotection^58^. Our *in vitro* results demonstrate that the present model is suitable for quantitative detection of UVB-dependent DNA photodamage, and show that natural and intact MCs, and so physiologically relevant MC^+^ organelles, are photoprotective structures requiring perinuclear positioning for optimal UVB protection (**Fig. 6, Extended Data Fig. 6**). Given that IF cages surround MC^+^ organelles regardless of the skin color type (**Fig. 3, Extended Data Fig. 3**), heterologous models could be generated in the future to test, for example, the photoprotective capacity of lightly or heavily melanized MCs in HEKs from donors with darker or lighter skin respectively, or to define the cellular/molecular mechanism underlying the distinct packaging of MCs in their organelles as a function of skin color ^11^. To conclude, beyond providing a physiologically relevant system allowing the *in vitro* study of photoprotection, we show that the cytoskeletons-driven 3D-position of MC^+^ organelles is key for their function. This result therefore predicts that disruption of the perinuclear distribution of pigments in skin keratinocytes would affect natural skin photoprotection.

Diseases characterized by abnormal localization of pigments in keratinocytes could target KRT5 and IFs and lead to more susceptibility in patients to develop skin cancers. Interestingly, the *KRT5* gene mutation characterized in DDD manifests as hyperpigmentation and other symptoms in patients ^32–34^. Histological examinations of the skin of these DDD patients show an altered distribution of melanin in basal keratinocytes ^32^. Since the same mutation introduced in the human keratinocytic cell line altered the elastic properties of the cytosol, which could cause MC^+^ organelles to disperse (**Fig. 5**, **Extended Data Fig. 5**), we hypothesize that KRT5^+^ IFs and their ability to constrain pigments in the perinuclear region are one of the targeted cellular processes in DDD. Interestingly, some DDD patients develop squamous cell carcinomas in the hyperpigmented area ^75,76^. This would suggest that disruption of the intracellular 3D-position of pigments induced by KRT5 could lead to an increased susceptibility to develop skin cancers due to suboptimal photoprotection. Future investigations exploiting the newly established model, combined with studies on patients with pigmentary skin disorders, would identify additional cellular and molecular mechanisms of keratinocytes contributing to both human skin pigmentation and photoprotection.

## Author contributions

S.B-M., L. S. and C.De. conceived the study; E.M., M.B., J-B.M., G.R. and C.De. contributed to fund the project; S.B-M., L. S., N.L., C.G., M.P., C.N-M., M.B., J-B.M., C.Du., F.B., G.R. and C.De. designed the research; S.B-M., L.S., A-S. M., N.L., V.F., J.S-C., R.A.J., M.R., C.G., M.P., I.H. and C.N-M. developed protocols; S.B-M., L.S., A-S. M., N.L., V.F., M.R., C.G., M.P., I.H. and C.N-M. performed the experiments; S.B-M., L.S., N.L., M.R., I.H., and C. De. collected the data; S.B-M., L.S., A-S. M., N.L., J-B.M. and C.De. analyzed the data; All authors provided intellectual support and interpreted the results; S.B-M. and C.De. wrote the initial manuscript with contributions from L.S., A-S. M., C.G., M.B., J-B.M., C.Du., F.B. and G.R.. L. S. and C. De. designed, prepared and edited the revised version of the manuscript. All authors contributed intellectual capital into the study and edited versions of the manuscript.

## Acknowledgments

This work was supported by the Institut National de la Santé et de la Recherche Médicale (INSERM), Institut Curie, Centre National de la Recherche Scientifique (CNRS), Agence Nationale de la Recherche ANR “MYOACTIONS” (ANR-17-CE11-0029-03 to Anne Houdusse and C.De.), “CILIOPHAGING” (ANR-22-CE14-0019-01 to E.M) and “MOBIDIC” (ANR-23-CE14-0041-02 to C.De. and E.M.), Fondation ARC pour la recherche sur le cancer (ARCPJA22020060002267 to C.De.), Fondation pour la Recherche Médicale (FRM EQU201903007827 Team label to G.R, SPF201909009097 to J.S-C., FDT202001010801 to S.B-M.), National Institutes of Health grants (R01 EY015625 to Michael S. Marks and G.R.), Biomolecular Analyses for Tailored Medicine in AcneiNversa (BATMAN) project (funded by ERA PerMed [JTC_2018] to M.B.). This work was also supported by the French National Research Agency through the “Investments for the Future” program (France-BioImaging, ANR-10-INSB-04) and we acknowledge the PICT-IBiSA, member of the France-BioImaging national research infrastructure supported by the CelTisPhyBio Labex (N° ANR-10-LBX-0038) part of the IDEX PSL (N°ANR-10-IDEX-0001-02 PSL), and the Necker SFR technical imaging platform. N.L. received funding from University Paris Saclay through a Contrat Doctoral Spécifique pour Normaliens (CDSN) from Ecole Normale Supérieure Paris-Saclay. S.B-M. received funding from the European Union’s Horizon 2020 Research and Innovation Programme under the Marie Sklodowska—Curie grant agreement No.666003 (Marie Sklodowska—Curie Actions, Institut Curie 3-I PhD Programme, IC-3i-PhD, grant agreement No.666003). This publication reflects only the author’s view and the European Research Agency is not responsible for any use that may be made of the information it contains.

We thank all the members of the laboratories as well as Pr. Andres Alcover and Dr. Nathalie Sauvonnet (Institut Pasteur, INSERM U1221 and Biomaterials and microfluidics platform, Paris, France), Dr. Florence Niedergang (Institut Cochin, INSERM U1016, Paris, France), and Dr. Franck Perez (Institut Curie, CNRS UMR144, Paris, France) for insightful discussions during the project; Dr. Carlos Kikuti (Institut Curie, CNRS UMR144, Paris, France) for discussions during the analysis of MCs-enriched fractions; Dr. Hugo Moreiras and Dr. Duarte C. Barral (NOVA Medical School, Faculdade de Ciências Médicas, Universidade NOVA de Lisboa, Lisboa, Portugal) for the sharing of initial protocols for MCs isolation; Dr. Fatemeh Rajabi (Gustave Roussy Cancer Center, INSERM U981, Villejuif, France) for sharing anti-CPD antibody; Dr. Eric Rubinstein (Sorbonne Université, INSERM, CNRS, CIMI-Paris, Paris, France) for sharing anti-CD63 antibody; Dr. Renata Basto (Institut Curie, CNRS UMR144, Paris, France) for sharing α-tubulin antibody; Prof. Rudolf Leube and Dr. Nicole Schwarz (Institute of Molecular and Cellular Anatomy, RWTH Aachen University, Aachen, Germany) for sharing KRT5-mCh plasmid.

## METHODS

### Cell culture

All cells were grown at 37°C under 5% CO_2_.

#### Primary cells

Normal human epidermal keratinocytes (HEKs) from highly or lightly pigmented skin donors were purchased from CellSystems, Sterlab, ATCC, or donated by L’Oréal Research and Innovation (isolated from foreskin of skin donor). Normal human epidermal melanocytes (HEMs) and human pigmented skins from light and dark skin donors ^11^ were donated by L’Oréal Research and Innovation (isolated from foreskin of dark skin donor). Human cells and skins were obtained from surgical residues after written informed consent from the donors according to the principles expressed in the Declaration of Helsinki and in article L.1243-4 of the French Public Health Code. This legislation does not require prior authorization by an ethics committee for sampling or use of surgical waste. HEKs were maintained undifferentiated in culture by passage at ∼75% confluency by gentle detachment via two sequential incubations using just enough volume to cover the cells: first with Versene solution [Thermo Fisher Scientific; 10 min, room temperature (RT)] and second with StemPro™ Accutase™ Cell Dissociation Reagent (Thermo Fisher Scientific; 15 min; 37°C). HEKs were used until passage five and maintained in culture in DermaLife Basal Medium (CellSystems) supplemented with DermaLife K Life (keratinocytes-supplemented medium; CellSystems) or in Dermal Cell Basal Medium (ATCC) supplemented with Keratinocyte Growth Kit (ATCC).

#### Cell lines

MNT-1 cells, originally obtained from Pr. Michael S. Marks (Children’s Hospital of Philadelphia, Philadelphia, USA), were cultured in DMEM supplemented with 20% v/v Fetal Bovine Serum (FBS), 10% (v/v) AIM-V medium, sodium pyruvate (1%, v/v), non-essential amino acids (1%, v/v), and penicillin-streptomycin (1%, v/v) (Gibco, Thermo Fisher Scientific), tested free of mycoplasma contaminations. For MCs isolation, MNT-1 cells were seeded at ∼15% confluency in 150 cm^2^ flasks (Corning, NY) and then cultured for 1 week in DMEM supplemented with 10% FBS, 100 U/mL penicillin G and 100 mg/mL streptomycin sulfate (Gibco, Thermo Fisher Scientific). HaCaT cells were grown in low calcium DMEM (Thermo Fisher Scientific) with 10% Chelex-treated FBS (BIORAD) ^77^ and a mix of ampicillin and streptomycin (100 U/mL and 100 µg/mL, respectively). HaCaT cells were passed at 60-80% confluency to avoid terminal differentiation as described ^78^.

### Antibodies

The following antibodies were used for immunoblotting (IB), immunofluorescence microscopy (IFM), or immuno-labeling by EM (IEM).

#### Primary antibodies

mouse anti-HMB45 (recognizing PMEL^+^ fibrils onto which melanin deposits, and used here as a ‘melanocore marker’; clone HMB45; ab787; Abcam; 1:400 [IFM]); mouse anti-TYRP1 (TA99/Mel-5; American Type Culture Collection); 1:40 [IEM]); mouse anti-CD63 (IgG2b monoclonal antibody generated and kindly provided by Dr. Eric Rubinstein ^79^ [Centre d’Immunologie et des Maladies Infectieuses, Paris, France]; 1:100 [IFM]); mouse anti-CPDs (clone TDM-2; NM-DND-001; Cosmo Bio LTD; 1:100 [IFM]); rabbit anti-LAMP1 (PA1-654A; Invitrogen, Thermo Fisher Scientific; 1:200 [IFM]); rabbit anti-KRT5 (EP1601Y; ab52635; Abcam; 1:400 [IFM]; 1:1000 [IB]); rabbit anti-KRT14 (clone Poly19053; BioLegend; 1:400 [IFM]); mouse anti-KRT14 (clone LL002; Abcam; 1:5000 [IB]); mouse anti-α-tubulin (T6199; Sigma-Aldrich; 1:200 [IFM]); rabbit anti-GAPDH (G9545; Sigma-Aldrich; 1:10000 [IB]); sheep anti-TGN46 (AHP500GT; Bio-Rad; 1:200 [IFM]); chicken anti-vimentin (ab24525; Abcam; 1:400 [IFM]; 1:10000 [IB]); mouse anti-Plectin (clone 10F6; Santa Cruz Biotechnology; 1:100 [IFM]).

#### Secondary antibodies

HRP-conjugated goat anti–mouse, and –rabbit (ab6721 and ab6789, respectively; Abcam; 1:10000 [IB]); Alexa Fluor (AF)-488, -555 or -647-conjugated anti-rabbit, anti-sheep, or anti-mouse, and fluorescent phalloidin (Invitrogen, Thermo Fisher Scientific; 1:200 [IFM]); rabbit anti-mouse (Z0412; Dako; 1:200 [IEM]).

### siRNAs/plasmid transfection

For small interfering RNA (siRNA) transfections, HEKs seeded on glass coverslips in 6-well tissue plates (4×10^4^ cells/well at day 0) were transfected twice at days 1 and 4 with respective siRNAs (0.2 μM) using Oligofectamine (Invitrogen, Thermo Fisher Scientific) according to manufacturer’s instructions. HEKs were collected at day 6 for IB (**Fig. 4a** and **Extended Data 4a** and **4g**) or MC-uptake assays (**Fig. 4d-f**, **Fig. 6c**, and **Extended Data Fig. 4d, 4j**, and **6b**). The sense strand of the following siRNAs was synthesized by Qiagen: Control (siCtrl): 5′-AATTCTCCGAACGTGTCACGT-3′; Keratin5 (siKRT5; four individual siRNAs targeting KRT5): siKRT5#1, 5′-AAGCAGTGTTTCCTCTGGATA-3′; siKRT5#2, 5′-TGCACTGATGGATGAGATTAA-3′; siKRT5#3, 5′-CGGAAGAGCTTCAAGAGCTAA-3′; siKRT5#4, 5′-CCTCCAGAATGTGTTCAATAA-3′; Vimentin (siVIM, two individual siRNAs): siVIM#1, 5′-CTGGCACGTCTTGACCTTGAA-3′; and siVIM#2, 5′-ATGGCTCGTCACCTTCGTGAA-3′; Keratin14 (siKRT14, four individual siRNAs): siKRT14#1, 5’-AACCTGCGCATGAGTGTGGAA-3’; siKRT14#2, 5’-CCAGGTGGGTGGAGATGTCAA-3’; siKRT14#3, 5’-CAGCGGCCTGCTGAGATCAAA-3’; siKRT14#4, 5’-CTGGCAATCAATACAGCTTCA-3’; Plectin1 (siPLEC1, four individual siRNAs): siPLEC1#1, 5’-CCAGACTAATATATTAATATA-3’; siPLEC1#2, 5’-CCGCCAGGTGAAGCTGGTGAA-3’; siPLEC1#3, 5’-CAAGGTGTACCGGCAGACCAA-3’; siPLEC1#4, 5’-CACAGTGGAGAAGATCATCAA-3’.

For plasmid transfection, WT or KRT5^DDD^ HaCaT cells, seeded on glass coverslips in 6-well tissue plates and allowed to internalize MC for 1 day, were transfected (50% confluency) with 2 μg of KRT5-mCherry encoding plasmid, using jetPRIME (Polyplus, Sartorius) according to manufacturer’s instructions. Cells were imaged maximum 24h after transfection.

### Base editing of HaCaT cells

*KRT5* mutations associated with Dowling-Degos Disease (KRT5^DDD^) were identified in the literature and led to 7 different mutations: two nonsense mutations, three frameshift mutations, and two mutations affecting the first ATG initiation codon. The KRT5 c.2T>C (p. Met1?) mutation found associated with DDD in a 4-generations Chinese family ^80^ was inserted in the HaCaT keratinocyte cell line ^81^. The small guide RNA (sgRNA) 5’-GAGACATGGTGGCTTGTTCC-3’ was designed using BE-Designer (http://www.rgenome.net/be-designer/) and sgRNA activity confirmed by Be-Hive (https://www.crisprbehive.design/single). The oligo encoding sgRNA and its reverse complement were purchased from Eurofins Genomics (Eurofins Genomics, Ebersberg, Germany). Sense and anti-sense oligos were heated and cooled down using a thermocycler for complementary annealing. Double-strand oligos were ligated into the pRG2 plasmid (Addgene, #104274) linearized by the BsaI restriction enzyme. 5×10^5^ HaCaT cells were transfected with 0.5 mg pRG2 and 1.5 mg pCMV_ABEmax using Lipofectamine 3000 (ThermoFisher Scientific), according to the manufacturer’s protocol. pCMV_ABEmax was a gift from David Liu (Addgene plasmid # 112095; http://n2t.net/addgene:112095; RRID:Addgene_112095) ^82^. Cells were collected after 5 days, and single-cell sorted based on their forward and side size scatters (Influx automatic cell-sorter; BD Bioscience, Le Pont de Claix, France). Selected clones were grown, and DNA was extracted for Sanger sequencing analysis. KRT5 c.2T>C homozygous clones and WT clones were further expanded to perform experiments.

### Reagents and drug treatment

HEKs that had internalized MCs for 1 day were incubated 4 h with DQ^TM^ Green BSA (Thermo Fisher Scientific, 50 μg/mL) and 30 min with LysoTracker^TM^ Red (Thermo Fisher Scientific, 50 nM) prior to analyses by FM (**Fig. 2g**). To disrupt microtubules ^83^, HEKs that had internalized MCs for 1 day were first incubated 90 min at 4°C, then 90 min at 37°C in presence of DMSO or 10 μM nocodazole (Sigma), prior to chemically fixation and IFM approach (**Fig. 4e-f**; **Extended Data Fig. 4n and 5b**) or UVB irradiation (**Fig. 6c**, **Extended Data Fig. 6b**).

### UVB exposure and imaging of UVB-dependent DNA marks

HEKs were grown at day 0 on glass coverslips in 24-well tissue plates (5×10^4^ cells/well; **Fig. 6a**) or, when siRNA treated, in 6-well tissue plates (2×10^4^ cells/well; **Fig. 6c**, **Extended Data Fig. 6b**), incubated or not with MCs (20 ng/μL) at day 1 (**Fig. 6a**), and exposed at day 2 with a single dose of UVB (312 nm; 0.5 or 1 J/cm^2^) using a Biosun machine (Vilber Lourmat, Suarlée, Belgium). In siRNA assays (**Fig. 6c, Extended Data Fig. 6b**), before UVB exposure, coverslips in 6-well tissue plates were transferred to 24-well plates. A Kodacel filter was placed on top of the culture plate to filter out any UVC and wavelength <290 nm. HEK medium was replaced by PBS before UVB exposure and immediately fixed with 4% (v/v) PFA in PBS at RT for 15 min. To image UVB-induced photolesions, fixed HEKs were then treated 30 min at 37°C with 2 M HCl to denature DNA before labeling with anti-CPD antibody.

### Immunoblotting

Cells were collected by Accutase-detachment, previously incubated with Versene, and followed by centrifugation (1,200 rpm, 5 min, 4°C), and the pellets were resuspended in cold lysis buffer [(20 mM Tris-HCl, 150 mM NaCl, 0.1% (v/v) Triton X-100, pH 7.2, supplemented with protease inhibitor cocktail (Roche), 20 min, 4°C]. Then, post-nuclear supernatants were obtained by centrifugation (10,000 rpm, 10 min, 4°C). The protein contents in cell lysates were determined using the Pierce™ BCA Protein Assay Kit (Thermo Fisher Scientific), and the concentrations were adjusted with sample buffer (250 mM Tris-HCl, 10% (v/v) SDS, 50% (v/v) Glycerol, 0.5 M β-mercaptoethanol, 0.5% (w/v) Bromophenol blue, pH 6.8) prior to samples boiling 5 min at 95°C. Equal amount of protein lysates were loaded on 4-12% Bis-Tris gels (NuPage, Invitrogen) and transferred to 0.2 μm pore-size nitrocellulose membranes (Millipore, GE Healthcare). Membranes were blocked 30 min in PBS supplemented with either 0.1% (v/v) Tween-20 and 4% (w/v) non-fat dried milk or 0.1% (v/v) Tween-20 and 5% (w/v) BSA and incubated up to 12 h at 4°C with respective primary antibodies, and 45 min with secondary antibodies diluted in PBS/ 0.1% Tween-20. Membranes were washed 30 min at least twice in PBS/ 0.1% Tween-20 and developed using ECL Plus Western blotting detection system (GE Healthcare) or Immobilon Western (Millipore) and visualized using Amersham Hyperfilm ECL (GE Healthcare). All molecular weights (MW) were in kDa.

### Melanocore isolation

MNT-1 cells (∼4×10^6^ cells) were seeded at ∼15% confluency and cultured in 150 cm^2^ flasks (Corning, NY) for 7 days. Conditioned medium was collected and centrifuged (300 *g*, 5 min) prior to be stored (-80°C) or filtered for MC isolation. To isolate MCs, conditioned medium was passed through porous filter-containing columns with a pore size of MWCO 300 kDa (Vivaspin Centricon; Sigma-Aldrich, Darmstadt, Germany) (**Extended Data Fig. 1a**, panel 1) ^15^. After four or five serial centrifugations (2663 *g*, 4°C, 60 min), the medium was passed through the filter, while the black MC-enriched fraction was retained in the pocket on top of the filter (**Extended Data Fig. 1a**, panels 2-3). The dark solution was then collected, transferred to a 1.5 mL tube, and centrifuged (10,000 rpm, 4°C, 30 min) until obtaining a dark pellet that was washed 5 min in PBS before being pelleted again by centrifugation (**Extended Data Fig. 1a**, panel 4) and resuspended in PBS to get the MCs-enriched fraction. Then, the melanin concentration was estimated, and the MC-enriched fraction was aliquoted, snap-frozen in liquid nitrogen and stored at -80°C or freshly used for experiments.

### Melanin estimation

The melanin concentration of each MCs-enriched fraction was estimated by absorption spectrophotometry (490 nm) ^5^ using a standard range of a reference solution with known concentrations of synthetic melanin (Sigma-Aldrich). The average melanin concentration among 43 different MCs-enriched fractions (**Extended Data Fig. 1b**) were adjusted to a working concentration of 3 ng/μL when incubated with cells (unless stated otherwise).

### Melanosome purification

Melanosome purification was performed as described in ^51,84^. Briefly, MNT-1 cells seeded in 10-cm dishes were grown to 90% confluence, collected by trypsinization, pelleted, and suspended in melanosome buffer (25 mM Hepes, 1 mM EDTA, 0.1 M EGTA, and 0.02% sodium azide, pH 7.4) supplemented with 0.25 M sucrose, and homogenized using a Dounce homogenizer. Lysates were centrifuged (10 min, 600 *g*, 4°C), and the supernatant was collected and loaded on 2 M sucrose diluted in melanosome buffer and centrifuged 30 min (11,000 *g*, 4°C). The melanosome-enriched fraction corresponding to a black top ring was carefully recovered and centrifuged 60 min (100,000 *g*, 4°C). Pelleted melanosomes were resuspended in PBS, prior to being processed (IFM, IEM; **Fig. 1b** and **1d**).

### Melanocores uptake assay

MCs-enriched fraction was diluted in HEK culture medium (3 ng/μL) and 250 μL were deposited on top of HEKs grown on glass coverslips before centrifugation (170 *g*, 5 min, RT) to allow the synchronous deposition of MCs on cells (**Fig. 2a**, **Extended Data Fig. 2a**). Then, cells were maintained at 37°C in 5% CO_2_ incubator and chemically fixed at different time points before analyses (IFM, EM). Note that time 0 corresponds to cells fixed after centrifugation (**Fig. 2a**, left panel).

### Immunofluorescence and fluorescence microscopy

#### Immunofluorescence

Cell monolayers seeded on glass coverslips were fixed 15 min at RT in PBS/ 4% PFA (v/v), washed three times in PBS and incubated 10 min in PBS/ 50 mM glycine. Coverslips were washed once in IF buffer (PBS/ 0.2% (w/v) BSA/ 0.1% (w/v) saponin), incubated with primary antibodies (1 h, RT) diluted in IF buffer, washed 3 times in IF buffer, incubated with secondary antibodies or phalloidin (30 min, RT) diluted in IF buffer, and washed 3 times in IF buffer and then PBS before mounting the coverslips onto glass slides using ProLong™ Gold Antifade Mount with DAPI (Thermo Fisher Scientific).

#### Epifluorescence or Confocal microscopy

Images were acquired on a laser scanning confocal microscope (Nikon AX) equipped with motorized stage and a CFI PlanApo Lambda D 60X OIL WD 0,15mm objective, with numerical aperture of 1,42 or a TIRF video-microscope (Nikon) (SFR imaging platform at INEM) or a an upright Leica DMI-5000B motorized stage with a 100X HCX PlanApo objective, with numerical aperture of 1.4, equipped with a CoolSNAP HQ2 Photometrics camera CCD cooled, and a metal-halide Leica EL 6000 lamp, and Metamorph, or a 3D Deconvolution widefield microscope (Ni-E; Nikon), equipped with a CoolSnap HQ2 camera (Photometrics) and a 100X Plan Apo VC oil objective. Z images series were acquired every 0.2 μm using the internal z-motor of the microscope with MetaMorph software (Molecular Devices) (PICT-IBiSA imaging platform at Institut Curie, Paris). Images were deconvolved with Meinel algorithm.

#### Super-resolution (SR) microscopy

images were acquired on a spinning-disk system based on an inverted Eclipse microscope (Ti-E; Nikon) integrated in Metamorph software (Molecular Devices, version 7.8.13), and equipped with a sCMOS camera (Prime 95B; Photometrics), a confocal spinning head (CSU-X1; Yokogawa), a 100x 1.4 NA Plan-Apo objective lens, and a super-resolution module (Live-SR; Gataca systems, France) based on structured illumination with optical reassignment technique and online processing leading to a two-time resolution improvement ^85^. The method, called multifocal structured illumination microscopy ^86^, allows combining the doubling resolution together with the physical optical sectioning of confocal microscopy. The maximum resolution is 128 nm with a pixel size in SR mode of 64 nm (**Fig. 2e**, **Extended Data Fig. 2e**).

#### Live cell microscopy

WT or KRT5^DDD^ HaCaTcells were grown on FluoroDish (WPI), incubated (30 min, 37°C) with SiR-tubulin (Spirochrome, 100 nM) to label MTs, washed once in culture medium and imaged at 37°C in their media supplemented with 20 mM HEPES (GIBCO, Thermo Fischer Scientific). Cells were imaged on a spinning disk CSU-X1 (Yokogawa) integrated in Metamorph software (Molecular Devices, version 7.8.13) with an EMCCD camera (iXon 897; Andor Technology), a 100× 1.4 NA Plan-Apo objective lens, and Z-images were taken at 0.2 μm with the piezoelectric motor (Nano z100, Mad City Lab). The analysis of individual MC^+^ organelle dynamics were performed using ImageJ FIJI (plugin MTrackJ) ^87^. Instantaneous speed was defined as displacement between two frames.

### Electron microscopy

Electron micrographs were acquired using a Tecnai Spirit G2 transmission electron microscope (FEI, Eindhoven, The Netherlands) operated at 80 kV and equipped with a 4k CCD camera (Quemesa, Olympus, Münster, Germany).

#### Conventional EM on ultrathin section

HEKs seeded on glass coverslips and allowed to internalize MCs were chemically fixed 24 h in 2.5% (v/v) glutaraldehyde in 0.1 M cacodylate buffer (pH 7,4), post-fixed 45 min in the dark (4°C) with 1% (w/v) osmium tetroxide supplemented with 1.5% (w/v) potassium ferrocyanide, dehydrated in ethanol and embedded in Epon as described in ^88^ (**Fig. 2c**, **Fig. 3a**, **Extended Data Fig. 2c** and **Extended Data Fig. 3a-b**). Ultrathin sections of 60-70 nm thickness were prepared with a Reichert UltracutS ultramicrotome (Leica Microsystems), post-stained with 4% aqueous uranyl acetate (10 min, RT) and lead citrate (1 min, RT). Conventional EM on human skin samples were performed as previously described ^11^ (**Extended Data Fig. 3a-b**).

#### Conventional EM on melanin particles

Fractions enriched in MCs or melanosomes (∼10 ng each) were deposited and allowed to settle for 30 min (RT) on copper/palladium Formvar coated and heavily carbonated grids. Excess of PBS was removed with Whatman filter paper without turning the grid and substituted by 5 μL of 2% (v/v) PFA in Phosphate Buffer (0,2 M, pH 7,4, RT, 20 min). Grids were extensively washed with distilled H_2_O, and samples were negatively stained 10-12 min at 4°C in the dark with a solution of uranyl acetate (UA, 4% in water) and methylcellulose (MTC, 2%) (UA/MTC; 1:9) (**Fig. 1a**, **Extended Data Fig. 1c**).

#### Immuno-EM on melanin particles

MCs or melanosomes were deposited on EM grids (see above) and fixed with 2% PFA in 0.1 M PB (pH 7.4) and processed for single immunogold labelling using anti-TYRP1 antibody (1 h, RT) followed by incubations with a rabbit anti-mouse antibody (bridge step, 20 min RT) and protein-A conjugated to 10 nm gold particles (Cell Microscopy Center, Department of Cell Biology, Utrecht University; 20 min, RT) (**Fig. 1d**).

### Optical Tweezers-Based Intracellular Micro-rheology

The setup combining optical tweezers and fast confocal microscopy was described in detail previously ^55^. Briefly, a single fixed optical trap was built by coupling a 1060-1100 nm infrared laser beam (2 W maximal output power; IPG Photonics) to the back port of an inverted Eclipse microscope (Nikon) equipped with a resonant laser confocal A1R scanner (Nikon), a 37°C incubator, and a nanometric piezostage (Mad City Labs).

WT and KRT5^DDD^ HaCaT cells were cultured on glass bottom dishes (Matek) and incubated with 2 µm diameter fluorescent beads (Invitrogen) for 24-48 h. Cells containing typically one to three internalized beads located in the perinuclear region were selected for the experiment. Images were taken in the resonant mode of the A1R scanner at 15.4 frames per second using a 100X Plan Apochromat objective with Zoom 4 giving a pixel size of 0.06215 µm/pixel.

Step relaxation experiments were described previously ^55^. Briefly, a 2 µm-diameter bead was first trapped in the optical tweezers (1 W laser output power, corresponding to 150 mW on the sample, trap stiffness *k_trap_* = 240 ± 40 pN.µm^-1^) calibrated by the Stokes drag force method). Following a *X_s_* = 0.5 μm step (linear increase in 40 ms) displacement of the stage, the bead was moved out of the trap center due to the viscoelasticity of its microenvironment. As the optical tweezers act as a spring on the bead, the bead position *x_b_*(*t*) relaxes from its maximal position *X_b_*, termed bead step amplitude, toward the center of the optical trap. Relaxation curves *x_b_*(*t*) during a duration *T* = 10 s are obtained with a home-made Matlab tracking code and analyzed with a phenomenological approach and a viscoelastic standard linear liquid (SLL) model as described previously ^55,89^. The phenomenological approach gives the bead step amplitude *X_b_* and the rigidity index *RI* defined as 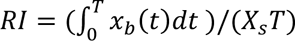 so that *RI* → 0 corresponds to a very soft and deformable cytoplasm while *RI* → 1 corresponds to a very stiff and rigid cytoplasm. The SLL model yields the spring constant (which corresponds to the elasticity) *K* (in N/m) and the viscosity *η* (in Pa.s) (**Fig. 5**, **Extended Data Fig. 5**).

### Image analysis and quantifications

#### Automatic detection of MCs and analysis

MCs incubated or not with HEKs are imaged over time (hours to days) by brightfield microscopy and/or fluorescence microscopy by immunolabeling using HMB45 antibody ^5^. The following codes were developed on Image J/Fiji software ^90^ to automatically detect and analyze MCs and MCs^+^ organelles across z-stack images.

#### Code_1_: Automatic detection of MCs, their spatial coordinates and analysis of associated fluorescence

Code_1_ combined two sequential steps: the generation of a region of interest (ROI) and the ROI analysis. *ROI generation:* First, MCs are automatically detected in a delimited region of interest (ROI). To detect MCs per cell or those deposited on coverslips, out-of-focus planes were removed, and then the cell contours or area of interest were manually drawn using the “freehand selection” tool and added to the ROI manager before running the macro. Note that when MCs were deposited on glass coverslips, a box corresponding to the size of the image was drawn using the “rectangle” tool. Then, a first detection of MCs is run automatically and separately for each predefined contour in the brightfield channel (**Extended Data Fig. 1e**, top panel; and **Fig. 2a**, middle insets) by creating a MIN-intensity z-projection and considering the lowest values (black spots), defined using the ‘Find Maxima’ function of Image J/Fiji, as the MCs. This was followed by a manual correction step to delete false-positive detections due to image noise or background. For each brightfield-detected spot, the focal z-plane for which the pixel intensity reached the minimum value (MCs being black at the focal plane) was defined prior to an additional (and optional) detection of MCs in the (red) fluorescence channel by creating a MAX-intensity z-projection and considering the highest values (red spots) as HMB45^+^ MCs. For each HMB45^+^ spot, the focal z-plane for which the pixel intensity reached the maximum value was defined. Automatic detection in the brightfield and fluorescence channels allowed to include in the analysis highly-pigmented HMB45^-^ MCs and low-pigmented HMB45^+^ MCs (**Extended Data Fig. 1e**, bottom panel, arrowheads; and **Extended Data Fig. 1f**). Automatically detected MCs were defined as ellipsoids centered on the spatial coordinates provided by the ‘Find Maxima’ plugin and whose radius (here, 1 µm in XY and in Z) can be adjusted by the user. Code_1_ generates a zip-file including the detected ROIs for MCs saved as circles located on the Z-slice defined as the focal plane. This ROI was used for further analysis (see Code_2_). *<u>ROI analysis</u>:* This second step was run after MCs automatic detection and analyzed the positive pixels detected in additional fluorescent channels of the image and within the ROI of each MC. Intensity analyses were performed on the 3 dimensions of the ROIs defined by Code_1_. Above a threshold (defined by the Triangle automatic threshold method, calculated on the maximum intensity projection), MC^+^ organelles were considered positive for the marker of interest whose percentage was calculated. Quantifications were from at least three independent experiments. *<u>Code_2_: Analysis of the MC-nuclear membrane distance.</u>* Code_2_ estimated the distance from the automatically detected MCs (using ROI set defined by code_1_) to the nuclear membrane. First, a manual correction step was performed to delete MCs considered to be outside of the cell. Then, code_2_ measured the Euclidian distance *(D)* from their center to the center of mass of the cell nucleus defined manually by drawing the nucleus contour at its focal plane (visually determined by DAPI staining). The nuclear radius *(r)* is estimated by averaging its longest and shortest axes. The subtraction of *r* from *D* defined the distance *(d),* [*D* – *r* = *d*], corresponding to the estimated distance in μm from each MC^+^ organelle to the nuclear membrane (NM) (**Extended Data Fig. 2b**).*<u>Code_3_: analysis of MCs vertical distance to the middle plane of the nucleus</u>*. Spatial coordinates of MCs (x, y, z) were retrieved by running Code_3_, thus obtaining the z-slice in which each MC was initially detected (Z_MC_). The mid-plane of the nucleus of each cell (Z_Nmid_) was calculated by determining the positions of the z-slice corresponding to the lower and upper parts of the nucleus, using DAPI fluorescence staining, and then applying the formula [Z_Nmid_=(Z_up_–Z_low_)/2]. Then, Z_Nmid_ was subtracted from the position of the z-slice of each detected MC, for each given cell (Z_MC_–Z_Nmid_). Negative values correspond to MCs positioned below the middle plane of the nucleus, and positive values to MCs above the mid-plane of the nucleus. The (Z_MC_–Z_Nmid_) differences were multiplied by the z-step (0.2 µm) to obtain the approximative vertical distance to the nucleus mid-plane and to calculate the median distance per cell (**Fig. 4h**, **Extended Data Fig. 4f and 4t**).

#### Image processing for 3D surface rendering

The acquired 3D image data was processed and analyzed using the Imaris software (version 9.8.2, Oxford Instruments). Initially, the raw image files (.stk format) underwent a pre-processing step, where they were deconvolved using Metamorph software (version 7.10.3; Molecular Devices) to improve signal-to-noise ratio and resolution. Following deconvolution, the processed image stacks were imported into Imaris for advanced 3D visualization. For 3D reconstruction, the Maximum Intensity Projection (MIP) algorithm was employed to generate a comprehensive view of the sample’s volume. Subsequently, the Imaris surface rendering module was used to accurately render the objects of interest in the dataset. The surface reconstruction was further refined using the “Automatic Threshold” function, allowing the identification and segmentation of objects based on intensity gradients within the 3D space (**Fig. 3c, Video 1**).

#### Immunogold labelling quantification

The average number of gold particles per pigment (isolated MC or melanosome) was calculated by dividing the total number of gold particles associated with pigment by the total number of pigmented structures (**Fig. 1e**).

#### CPDs intensity quantification

HEKs with or without internalized MCs were UVB irradiated. Code_1_ was used to obtain the number of MCs/cell, then the contour of the nucleus of cells was manually drawn based on their respective DAPI staining and kept as ROIs. Cells were imaged from top to bottom with a z-step of 0.2 μm, and image stacks were z-projected (maximum-intensity). The mean fluorescence intensity of CPD staining/ROI was measured after background subtraction, and the effect on the photoprotection was defined by normalizing the CPDs intensities to the mean value in control cells (without internalized MCs). Fiji software was used for image analysis (**Fig. 6b** and **6d**).

#### Quantification of structures in the vicinity of MC^+^ organelles by conventional EM

Intracellular structures in the vicinity of MC^+^ organelles were quantified on 2D-EM images by depicting a 1 μm-diameter circle centered around each MC^+^ organelle and by measuring the number of organelles and other structures in each circle (**Extended Data Fig. 2d**). The 1 μm-diameter was defined approximatively as twice the average diameter of a MC^+^ organelle (∼500 nm), so that the subcellular environment considered would at least be ∼250 nm distant from the limiting membrane of each MC^+^ organelle (**Extended Data Fig. 2d**). Identification of intracellular organelles/structures was based on their morphology: i.e., mitochondria (M): double membrane-bound compartment with internal cristae; endoplasmic reticulum (ER): irregular membrane compartment decorated by electron dense cytosolically-exposed (ribosomal) densities; Golgi apparatus (GA): tubulovesicular structures associated with cisternae; intermediate filaments made of keratin bundles (K): electron dense ‘hairy’ cytosolic and filamentous structures; plasma membrane (PM): membrane separating the cell interior from its exterior; nuclear membrane (NM): membrane separating the nuclear material from the cytosol; Endo/lysosomal compartments (EL): membrane compartments with electron-lucent lumen filled with various amount of intraluminal vesicles and/or electron dense materials; tubulo-vesicles (TVL): tubulo-vesicular structures of unknown identity. The number of organelles identified was counted using ImageJ/Fiji and expressed as a percentage of the total number of MC^+^ organelles (**Fig. 2d**).

### Statistical analysis

Statistical analyses were performed using GraphPad Prism (version 7 and 8, GraphPad Software, San Diego California, USA, www.graphpad.com). Scored or quantified cells in each experiment were randomly selected, and all experiments were repeated at least three times. Results were reported as mean or median ± standard error of the mean (s.e.m.). Unless otherwise mentioned, statistical analyses between (i) two sets of data were performed by using the two-tailed unpaired Student’s t-test, (ii) more than two sets of data were performed with an ordinary one-way ANOVA with a Tukey post-hoc correction for multiple comparisons. In such cases, the reported *P* values are multiplicity adjusted *P* values for each comparison. In all statistical analysis, significant differences between control or treated samples are indicated as \*\*\*\**P*<0.0001, \*\*\**P*<0.001, \*\**P*<0.01, *P<0.05. Only *P*<0.05 was considered as statistically significant. Correlation analyses between multiple variables were performed using correlation matrix computing Spearman non-parametric correlation coefficient (r) for every pair of data sets and the associated two-tailed *P* values presented as heatmap of the correlation matrix. For detailed results, see Table 1.

**Table 1.**

| Figure/panel | Condition | Data | Number (n)/experiment |
| --- | --- | --- | --- |
| Fig. 1c | Melanocores<br>Melanosomes | 92.3 ± 1.7 %<br>98.9 ± 0.6 % | 511 MCs, 3 experiments<br>492 melanosomes, 3 experiments |
| Fig. 1e | Melanocores<br>Melanosomes | 2.7 ± 0.6<br>8.5 ± 1.7 | 29 MCs, 77 gold particles,<br>3 experiments<br>29 melanosomes, 246 gold particles,<br>3 experiments |
| Extended Data Fig. 1b | Melanin<br>concentration | 104 ± 17 ng/μL | 41 MC-enriched fractions |
| Fig. 2b | 10min<br>1h<br>8h<br>1day<br>3day<br>7day | 13.6 ± 0.6 μm<br>6.9 ± 0.7 μm<br>2.7 ± 0.4 μm<br>1.2 ± 0.17 μm<br>1.2 ± 0.13 μm<br>0.9 ± 0.07 μm | 41 MC<sup>+</sup> organelles, 6 cells,<br>3 experiments<br>47 MC<sup>+</sup> organelles, 5 cells,<br>3 experiments<br>39 MC<sup>+</sup> organelles, 5 cells,<br>3 experiments<br>103 MC<sup>+</sup> organelles, 9 cells,<br>3 experiments<br>105 MC<sup>+</sup> organelles, 10 cells,<br>3 experiments<br>214 MC<sup>+</sup> organelles, 12 cells,<br>3 experiments |
| Fig. 2d | PM<br>GA<br>ER<br>Mito<br>NM<br>KRT<br>Endo<br>TVM | 6.1 %<br>9.1 %<br>18.2 %<br>27.3 %<br>42.4 %<br>50 %<br>53 %<br>66 % | 66 MC<sup>+</sup> organelles, 2 experiments |
| Fig. 2f | LAMP1<br>CD63 | 85.2 ± 7.2 %<br>97.1 ± 1.3 % | 475 MC<sup>+</sup> organelles, 36 cells,<br>3 experiments |
| Fig. 2h | Lysotracker<br>DQ-BSA | 24.1 ± 6.5 %<br>13 ± 2.8 % | 1186 MC<sup>+</sup> organelles, 68 cells,<br>3 experiments |
| Fig. 3d | KRT5<br>MT<br>F-actin | 45.4 ± 4.6 %<br>26.1 ± 4.8 %<br>14.3 ± 3.0 % | 248 MC<sup>+</sup> organelles, 24 cells,<br>3 experiments |
| Fig. 4b | siKRT5 | 21.4 ± 11.6 % | 3 experiments |
| Fig. 4c | siKRT5 | 27.6 ± 7.6 % | 77 cells, 3 experiments |
| Fig. 4g | siCTRL-DMSO<br>siCTRL-NOCO<br>siKRT5-DMSO<br>siKRT5-NOCO | 0.59 ± 0.05 μm<br>0.88 ± 0.09 μm<br>1.56 ± 0.11 μm<br>2.1 ± 0.16 μm | 2598 MC<sup>+</sup> organelles, 111 cells,<br>3 experiments<br>2186 MC<sup>+</sup> organelles, 64 cells,<br>3 experiments<br>1790 MC<sup>+</sup> organelles, 97 cells,<br>3 experiments<br>2486 MC<sup>+</sup> organelles, 80 cells,<br>3 experiments |
| Fig. 4h | siCTRL-DMSO<br>siCTRL-NOCO<br>siKRT5-DMSO<br>siKRT5-NOCO | 0.94 ± 0.22 μm<br>0.24 ± 0.11 μm<br>0.22 ± 0.14 μm<br>-0.08 ± 0.12 μm | 2643 MC<sup>+</sup> organelles, 43 cells,<br>3 experiments<br>2046 MC<sup>+</sup> organelles, 48 cells,<br>3 experiments<br>1930 MC<sup>+</sup> organelles, 42 cells,<br>3 experiments<br>2470 MC<sup>+</sup> organelles, 52 cells,<br>3 experiments |
| Extended Data Fig. 4b | KRT14 (siKRT5)<br>KRT14 (siKRT14)<br>KRT5 (siKRT5) | 87.4 ± 9.3 %<br>53.8 ± 14.3 %<br>25.0 ± 9.7 % | 3 experiments<br>3 experiments<br>3 experiments |
|  | KRT5 (siKRT14) | 86.3 ± 2.9 % | 3 experiments |
| Extended Data Fig. 4c | siKRT14 | 30.4 ± 12.9 % | 52 cells (siKRT14), 64 cells (siCTRL), 3 experiments |
| Extended Data Fig. 4e | siCTRL-DMSO | 0.26 ± 0.06 µm | 826 MC<sup>+</sup> organelles, 33 cells, 3 experiments |
|  | siKRT14-DMSO | 1.00 ± 0.18 µm | 1071 MC<sup>+</sup> organelles, 31 cells, 3 experiments |
| Extended Data Fig. 4f | siCTRL-DMSO | 0.47 ± 0.16 µm | 3556 MC<sup>+</sup> organelles, 80 cells, 3 experiments |
|  | siKRT14-DMSO | -0.15 ± 0.13 µm | 1137 MC<sup>+</sup> organelles, 33 cells, 3 experiments |
| Extended Data Fig. 4h | siVIM | 22.6 ± 7.9 % | 3 experiments |
| Extended Data Fig. 4i | siVIM | 13.4 ± 3.5 % | 63 cells, 3 experiments |
| Extended Data Fig. 4k | siCTRL | 0.73 ± 0.08 µm | 1522 MC<sup>+</sup> organelles, 44 cells, 3 experiments |
|  | siVIM | 0.53 ± 0.11 µm | 842 MC<sup>+</sup> organelles, 27 cells, 3 experiments |
| Extended Data Fig. 4l | siCTRL | 34.6 ± 4.7 | 1522 MC<sup>+</sup> organelles, 44 cells, 3 experiments |
|  | siVIM | 31.2 ± 3.4 | 842 MC<sup>+</sup> organelles, 27 cells, 3 experiments |
| Extended Data Fig. 4m | siCTRL | 6.7 ± 0.2 µm | 121 cells, 3 experiments |
|  | siVIM | 7.1 ± 0.3 µm | 44 cells, 3 experiments |
|  | siCTRL-DMSO | 7.2 ± 0.2 µm | 50 cells, 3 experiments |
|  | siCTRL-NOCO | 7.3 ± 0.4 µm | 39 cells, 3 experiments |
|  | siKRT5-DMSO | 7.4 ± 0.2 µm | 76 cells, 3 experiments |
|  | siKRT5-NOCO | 7.2 ± 0.2 µm | 33 cells, 3 experiments |
|  | siKRT14-DMSO | 6.8 ± 0.2 µm | 53 cells, 3 experiments |
|  | siPLEC1-DMSO | 6.9 ± 0.34 µm | 33 cells, 3 experiments |
| Extended Data Fig. 4p | siPLEC1 | 0.01 ± 0.003 | 3 experiments |
| Extended Data Fig. 4q | siPLEC1 | 0.30 ± 0.03 | 33 cells, 3 experiments |
| Extended Data Fig. 4s | siCTRL-DMSO | 1.08 ± 0.14 µm | 39 cells, 3 experiments |
|  | siPLEC1-DMSO | 0.98 ± 0.16 µm | 33 cells, 3 experiments |
| Extended Data Fig. 4t | siCTRL-DMSO | 0.14 ± 0.15 µm | 83 cells, 3 experiments |
|  | siPLEC1-DMSO | -1.22 ± 0.21 µm | 33 cells, 3 experiments |
| Fig. 5b | HaCaT WT |  | 34 beads, 4 experiments |
|  | HaCaT KRT5<sup>DDD</sup> |  | 27 beads, 4 experiments |
| Fig. 5c | HaCaT WT | 0.305 ± 0.059 µm | 34 beads, 4 experiments |
|  | HaCaT KRT5<sup>DDD</sup> | 0.247 ± 0.073 µm | 27 beads, 4 experiments |
| Fig. 5d | HaCaT WT | 0.321 ± 0.124 | 34 beads, 4 experiments |
|  | HaCaT KRT5<sup>DDD</sup> | 0.224 ± 0.132 | 27 beads, 4 experiments |
| Fig. 5e | HaCaT WT | 175.2 ± 102.1 pN/µm | 34 beads, 4 experiments |
|  | HaCaT KRT5<sup>DDD</sup> | 132.5 ± 101.6 pN/µm | 27 beads, 4 experiments |
| Extended Data Fig. 5b | HaCaT KRT5<sup>DDD</sup> + NOCO | 1.11 ± 0.14 µm | 1118 MC<sup>+</sup> organelles, 54 cells, 3 experiments |
|  | HaCaT KRT5<sup>DDD</sup> + NOCO + KRT5-mCh | 0.71 ± 0.10 µm | 1614 MC<sup>+</sup> organelles 67 cells, 3 experiments |
| Extended Data Fig. 5c | HaCaT WT |  | 17 tracked MC<sup>+</sup> organelles, 7 cells, 2 experiments |
|  | HaCaT KRT5<sup>DDD</sup> |  | 20 tracked MC<sup>+</sup> organelles, 6 cells, 2 experiments |
| Extended Data Fig. 5g | HaCaT WT | 19.91 ± 13.55 Pa.s | 34 beads, 4 experiments |
|  | HaCaT KRT5<sup>DDD</sup> | 15.34 ± 10.75 Pa.s | 27 beads, 4 experiments |
| Fig. 6b | -MCs, (0.5 J/cm<sup>2</sup>) | 100.0 ± 5.6 | 62 cells, 4 experiments |
|  | +MCs, (0.5 J/cm<sup>2</sup>) | 46.1 ± 5.2 | 66 cells, 4 experiments |
|  | -MCs, (1 J/cm<sup>2</sup>) | 100.0 ± 6.2 | 22 cells, 3 experiments |
|  | +MCs, (1 J/cm<sup>2</sup>) | 54.7 ± 10.8 | 23 cells, 3 experiments |
| Fig. 6d | siCTRL-DMSO | 100.0 ± 7.2 | 80 cells, 4 experiments |
| | siCTRL-NOCO | $135.1 \pm 15.2$ | 51 cells, 3 experiments |
| | siKRT5-DMSO | $101.8 \pm 13.9$ | 37 cells, 3 experiments |
| | siKRT5-NOCO | $209.9 \pm 14.7$ | 86 cells, 4 experiments |
| Fig. 6e | siCTRL-DMSO +<br>siKRT5-NOCO | Spearman correlation;<br>0.317, p-value: 0.0002 | 132 cells, 5 experiments |
| Extended Data Fig. 6a | 0.5 J/cm<sup>2</sup> | Spearman correlation;<br>-0.32, p-value: 0.0002 | 129 cells, 4 experiments |
|  | 1 J/cm<sup>2</sup> | Spearman correlation;<br>-0.32, p-value: 0.027 | 47 cells, 3 experiments |
| Extended Data Fig. 6c | siCTRL-DMSO | $56.0 \pm 4.0$ | 4481 MC<sup>+</sup> organelles, 80 cells,<br>4 experiments |
| | siCTRL-NOCO | $44.3 \pm 4.2$ | 2261 MC<sup>+</sup> organelles, 51 cells,<br>3 experiments |
| | siKRT5-DMSO | $45.7 \pm 4.4$ | 1689 MC<sup>+</sup> organelles, 37 cells,<br>3 experiments |
| | siKRT5-NOCO | $46.5 \pm 4.3$ | 3906 MC<sup>+</sup> organelles, 84 cells,<br>4 experiments |
| Extended Data Fig. 6d | siCTRL-DMSO +<br>siKRT5-NOCO | Spearman correlation;<br>-0.058, p-value: 0.361 | 252 cells,<br>7 experiments |

### Material availability

All unique/stable reagents generated in this study are available from the lead contact with a completed materials transfer agreement.

**Video 1. 3D organization of MC^+^ organelles in contact with KRT5^+^ intermediate filaments.**

Animation of the 3D surface rendering of the image shown in Figure 3c. MC^+^ organelles (labelled with HMB45 antibody, red) are distributed in close proximity to KRT5^+^ intermediate filaments (green). Semi-transparency of the green channel was applied to better appreciate proximity. Scale bar: 5 µm.

**Video 2. MC^+^ organelles are confined in KRT5-expressing keratinocyte cells.**

Spinning-disc confocal microscopy on HaCaT-WT cells that internalized MCs (not shown) for 1 day and incubated with SiR-Tubulin probe (magenta). Trajectories of MC^+^ organelles (colored lines) throughout >8min of live imaging acquisition are shown. Acquisition parameters: 1 image (merge)/ 7 sec. Video is shown at 15 frame/s. Bar: 15 µm. See also **Extended Data Fig. 5c**.

**Video 3: MC^+^ organelles are mobile in KRT5-null keratinocyte cells.**

Spinning-disc confocal microscopy on HaCaT-KRT5^DDD^ cells that internalized MCs (not shown) for 1 day and incubated with SiR-Tubulin probe (magenta). Trajectories of MC^+^ organelles (colored lines) throughout >8min of live imaging acquisition are shown. Acquisition parameters: 1 image (merge)/ 3 sec. Video is shown at 15 frame/s. Bar: 15 µm. See also **Extended Data Fig. 5c**.

## Extended Data Figure legends

**Extended Data Figure 1.**
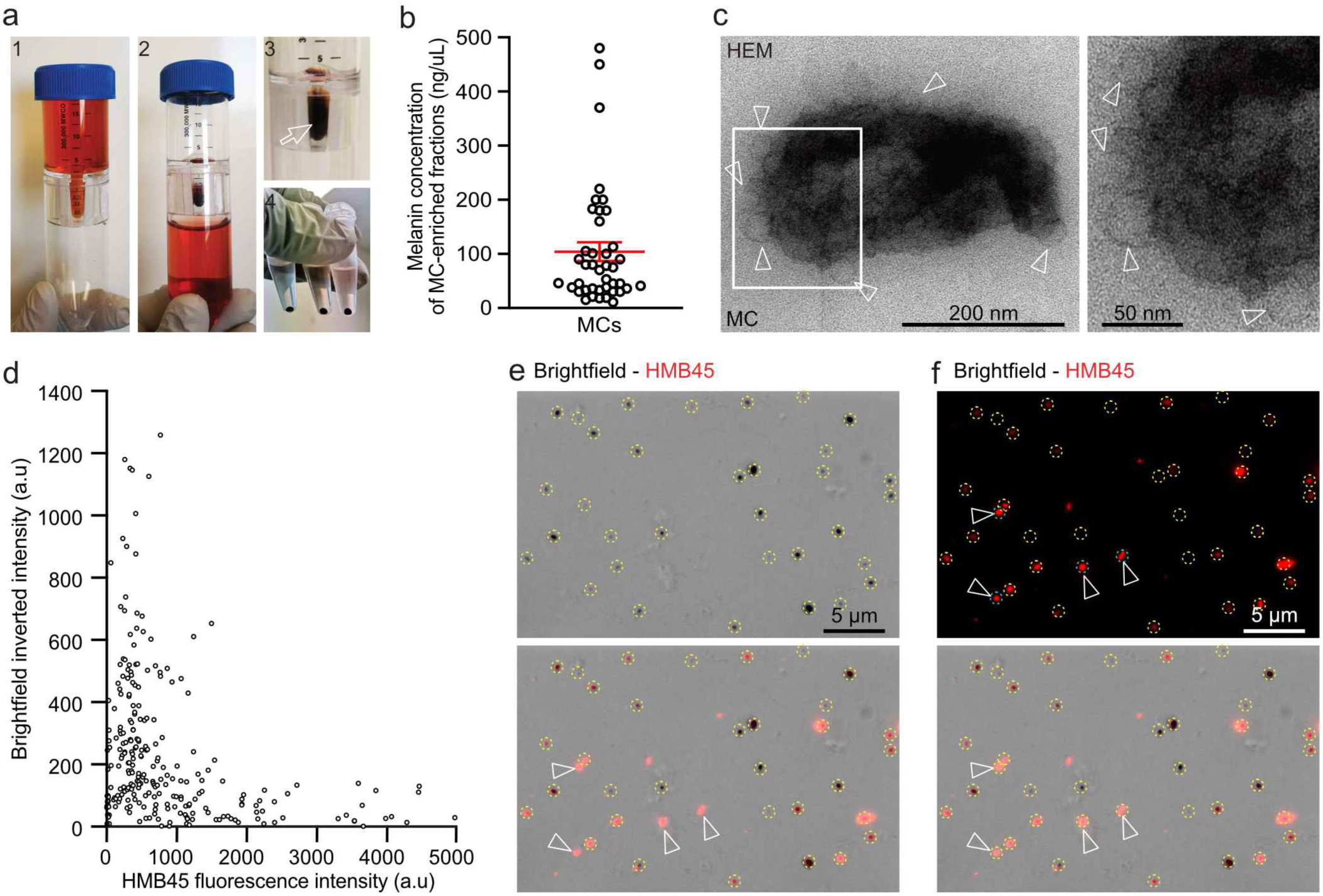
Collection and automatic optical detection of melanocores. (**a**) Steps illustrating the procedure for isolating MCs from MNT-1 cell culture medium collected and deposited in a column containing a porous filter (1), then centrifuged (2) to retain pigmented MCs in a pocket (3, arrow) followed by content transfer to a tube and centrifugation to form a pigmented MCs pellet (4). (**b**) Quantification of the average estimated melanin concentration of MCs-enriched fractions. (**c**) EM micrographs of MCs isolated from primary highly pigmented HEM cell culture medium. Arrowheads point to lipid vesicles associated to MC. (**d**) Quantification of the maximum intensity of the inverted signal captured by brightfield illumination of individual MCs as a function of their maximum intensity of the fluorescent signal for HMB45 (n=255 MCs). (**e**-**f**) IFM of isolated MCs captured by brightfield illumination (**e**, top) and stained with HMB45 antibody (**f**, top; red). HMB45^+^ and lighter MCs (**e**, bottom, arrowheads) not identified by automatic detection of MCs based on the brightfield image were subsequently identified using automatic detection of HMB45 signal (**f**, bottom, arrowheads). Automatically detected MCs are circled in yellow. Data are the average of at least three independent experiments presented as the mean ± SEM.

**Extended Data Figure 2.**
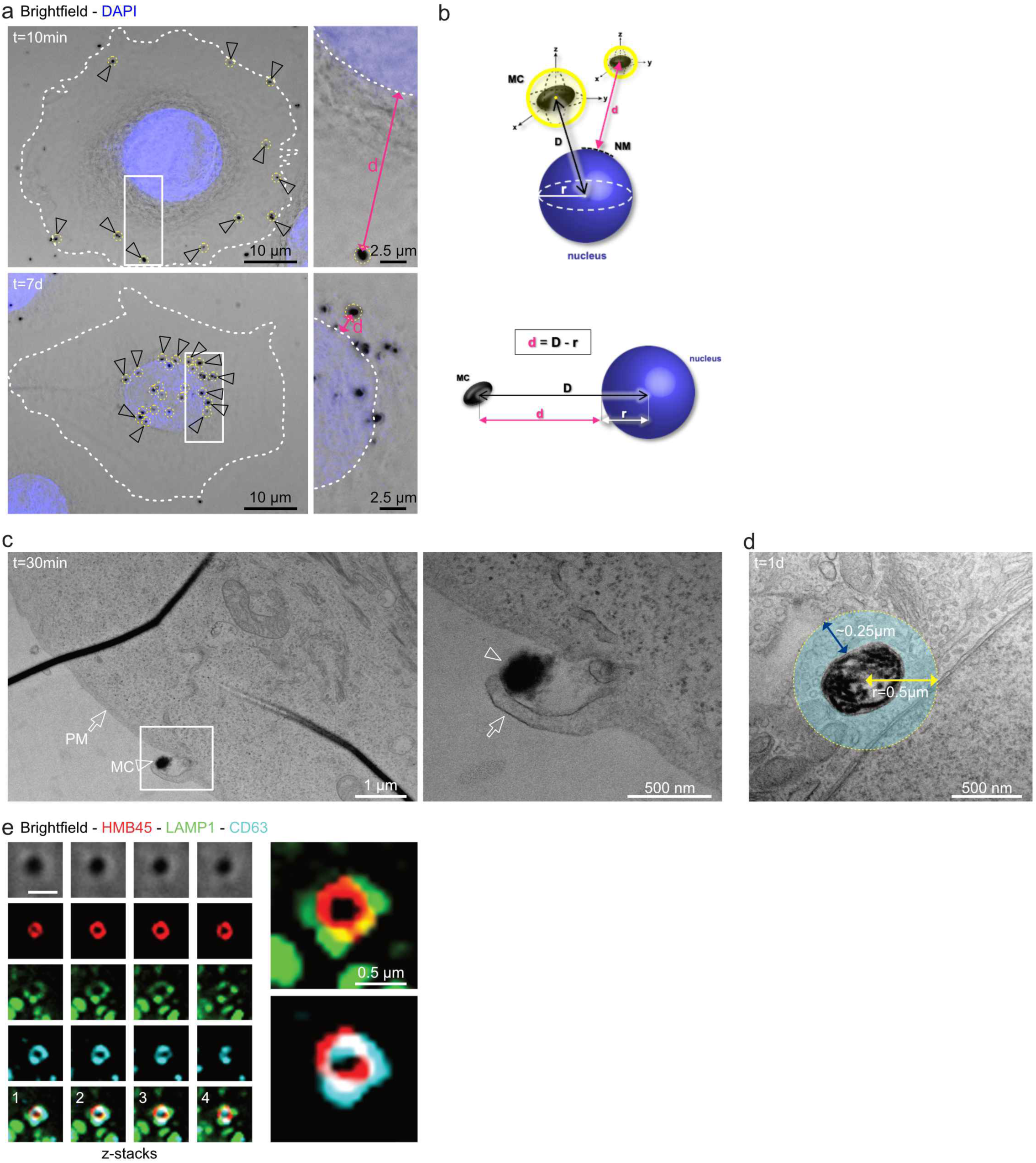
Position of melanocores organelles relative to nucleus and molecular composition. (**a**) IFM of HEKs 10 min (top) or 7-days (bottom) after MCs (arrowheads) deposition and DAPI staining (blue). Automatically detected MCs are circled in yellow and distance (d in insets) of individual MC to nucleus edge (dashed line in insets) is depicted in magenta. (**b**) Diagram showing the automatic measurement (by code_2_) of the total distance (D; double black arrow) from each MC (yellow circles) to the center of mass of the nucleus (blue ball). Further subtraction of the nuclear radius (r) from D gives the estimated 2D distance (d; magenta double arrow) from individual MC to the nuclear membrane (NM, dashed black line). (**c**) Conventional EM micrographs of ultrathin sections of HEKs 30 min after MC deposition. Arrowheads point to an extracellular MC affixed to plasma membrane (arrow) forming finger-like protrusion (inset, arrow). (**d**) Conventional EM micrograph of ultrathin section of HEKs 1-day after MC deposition depicting the area used for analysis in Fig. 2d. (**e**) SR-IFM of HEKs 1-day after MCs deposition and stained with HMB45 (red), anti-LAMP1 (green) and -CD63 (cyan) antibodies. Insets (left) are consecutive z-stacks of the same MC^+^ organelle.

**Extended Data Figure 3.**
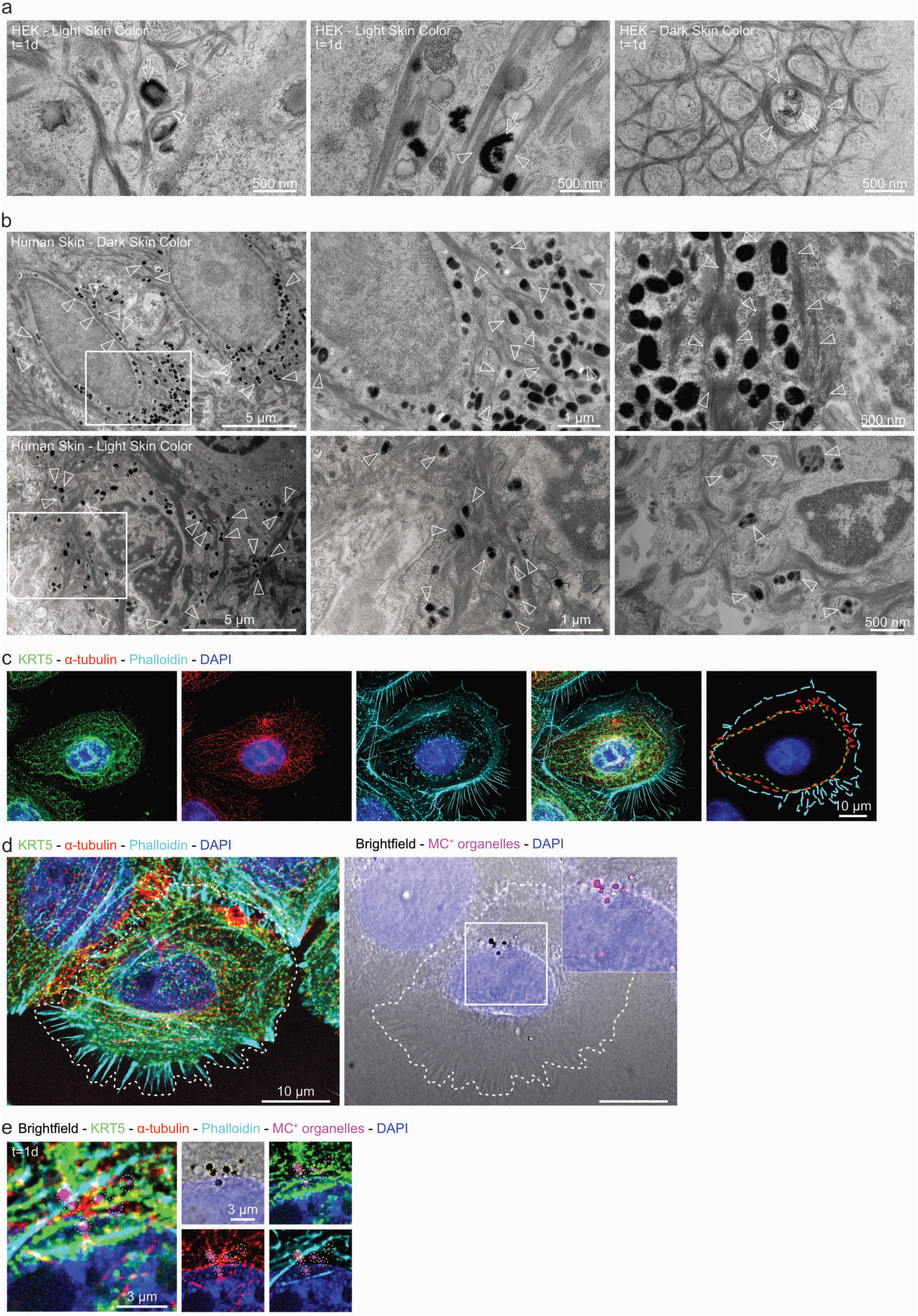
Melanocore organelles in keratinocytes are partially surrounded by keratin intermediate filaments in vitro and in vivo. (**a**) Conventional EM micrographs of ultrathin sections of HEKs from light (left, middle) or dark (right) skin donors 1 day after MC deposition. Arrowheads point to keratin^+^ intermediate filaments surrounding MC^+^ organelles (arrows). (**b**) Conventional EM micrographs of ultrathin sections of human pigmented skins from dark (top) or light (bottom) skin donors showing keratin^+^ intermediate filaments (arrowheads) surrounding MC^+^ organelles in epidermal keratinocytes. Middle panels are magnified area of the boxed regions. (**c**) IFM of HEKs stained with anti-KRT5 (green) and -α-tubulin (red) antibodies, fluorescence-conjugated phalloidin (cyan), and DAPI (blue). Monochrome and merged images are shown in addition to the outer contour delineation of the different cytoskeletal staining (dashed lines, right). (**d-e**) IFM of HEKs 1-day after MCs deposition (d) and stained as in (c) highlighting perinuclear MC^+^ organelles (captured by brightfield and pseudocolored in magenta) in. Automatically detected MCs are circled in yellow (e).

**Extended Data Figure 4.**
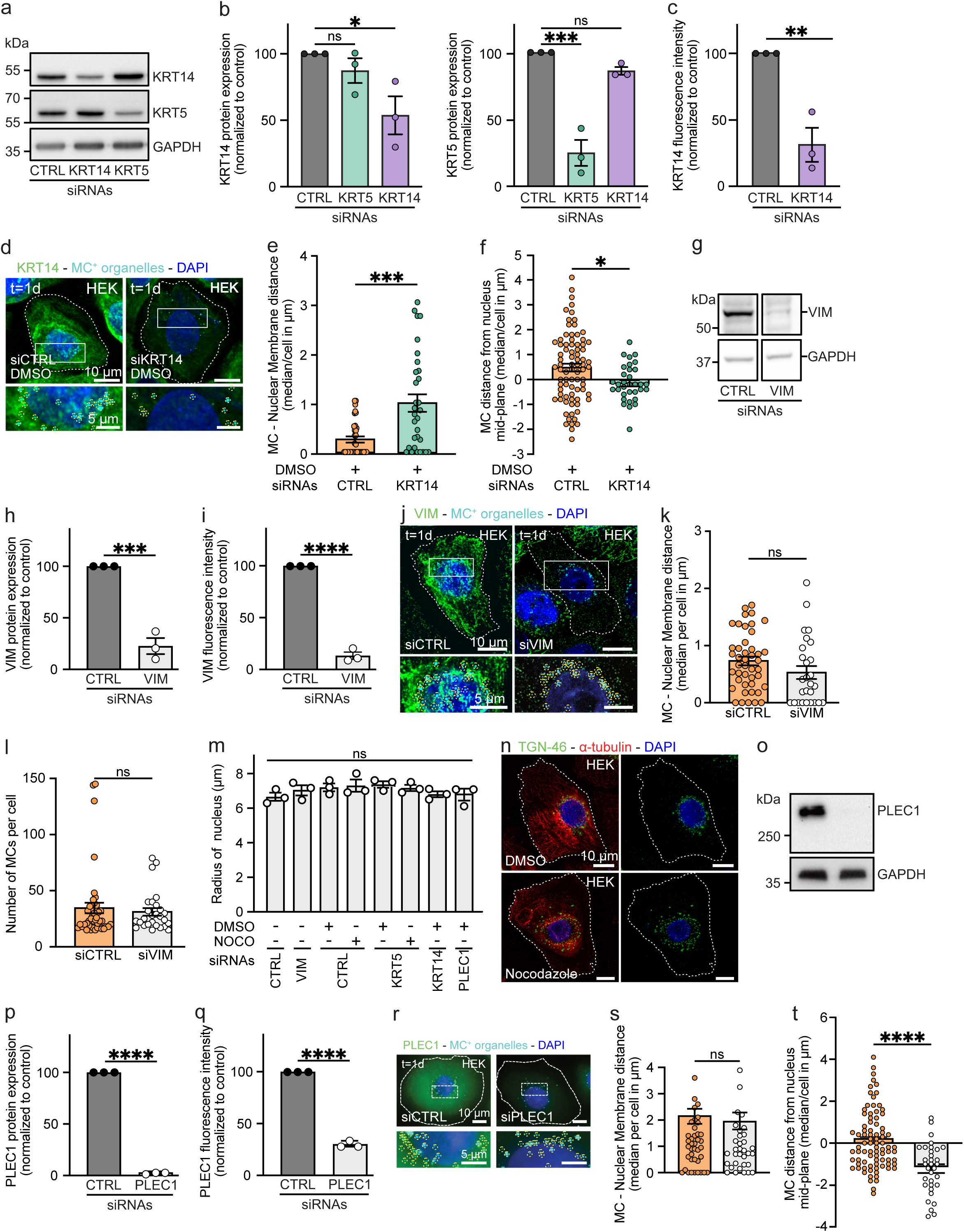
Keratin 14 controls the 3D-position of melanocore organelles, while plectin 1 contributes to their vertical positioning. (**a**) Western blot of HEK lysates treated with control (CTRL), keratin 14 (KRT14) or keratin 5 (KRT5) siRNAs and probed with anti-KRT14 (top), -KRT5 (middle) or -GAPDH (loading control, bottom) antibodies. (**b**) Quantification of the average expression levels of KRT14 (left) or KRT5 (right) in siRNA-treated HEKs (as in a) normalized to GAPDH levels and control. (**c**) Average fluorescence intensity of KRT14 in siCTRL- or siKRT14-treated HEKs analyzed in Fig. 4e-f and normalized to control. (**d**) IFM of siCTRL-(left) or siKRT14-(right) treated HEKs stained with anti-KRT14 antibody (green) and DAPI (blue). (**e**) Quantification of the median distance of MC^+^ organelles to the nucleus edge in siCTRL- or siKRT14-treated HEKs expressed per cell. (**f**) Quantification of the median distance of MC^+^ organelles from nucleus mid-plane expressed per cell. (**g**) Western blot of HEK lysates treated with CTRL or vimentin (VIM) siRNAs and probed for VIM (top) or GAPDH (loading control, bottom). (**h**) Quantification of the average expression level of VIM in siCTRL- or siVIM-treated HEKs normalized to GAPDH levels and control. (**i**) Quantification of the mean VIM fluorescence intensity in siCTRL- or siVIM-treated HEKs used for analysis in Extended Data Fig. 4j-k and normalized to control. (**j**) IFM of siCTRL-(left) or siVIM- (right) treated HEKs stained with anti-VIM antibody (green) and DAPI (blue). (**k**) Quantification of the median distance of MC^+^ organelles to the nucleus edge in siCTRL- or siVIM-treated HEKs expressed per experiment. (**l**) Quantification of the number of MC^+^ organelles in siCTRL- and siVIM-treated HEKs analyzed in (k). (**m**) Quantification of the mean radius of the nucleus in HEKs treated with CTRL, KRT5, KRT14, VIM or PLEC1 siRNAs and incubated or not with DMSO or nocodazole. (**n**) IFM of DMSO- or nocodazole-treated HEKs stained with anti-α-tubulin (red) and -TGN46 (green) antibodies, and DAPI (blue). (**o**) Western blot of HEK lysates treated with CTRL or plectin 1 (PLEC1) siRNAs and probed for anti-PLEC1 (top) or -GAPDH (loading control, bottom) antibodies. (**p**) Quantification of the average expression level of PLEC1 in siCTRL- or siPLEC1-treated HEKs normalized to GAPDH levels and control. (**q**) Quantification of the mean PLEC1 fluorescence intensity in siCTRL- or siPLEC1-treated HEKs used for analysis (Extended Data Fig. 4r-t) and normalized to control. (**r**) IFM of siCTRL- (left) or siPLEC1- (right) treated HEKs stained with anti-PLEC1 antibody (green) and DAPI (blue). (**s**) Quantification of the median distance of MC^+^ organelles to the nucleus edge in siCTRL- or siPLEC1-treated HEKs expressed per cell. (**t**) Quantification of the median distance of MC^+^ organelles from nucleus mid-plane expressed per cell. (d, j, n, r) Cell outlines are delineated by dashed lines. (d, j, r) Automatically detected MCs captured 1-day after deposition by brightfield microscopy were pseudo-colored in cyan and circled in yellow. Data are the average of at least three independent experiments presented as the mean or median ± SEM. Two-tailed unpaired t test and one-way ANOVA with Tukey post-hoc (ns, non-significant; **, *P* < 0.01; ***, *P* < 0.001; ****, *P* < 0.0001).

**Extended Data Figure 5.**
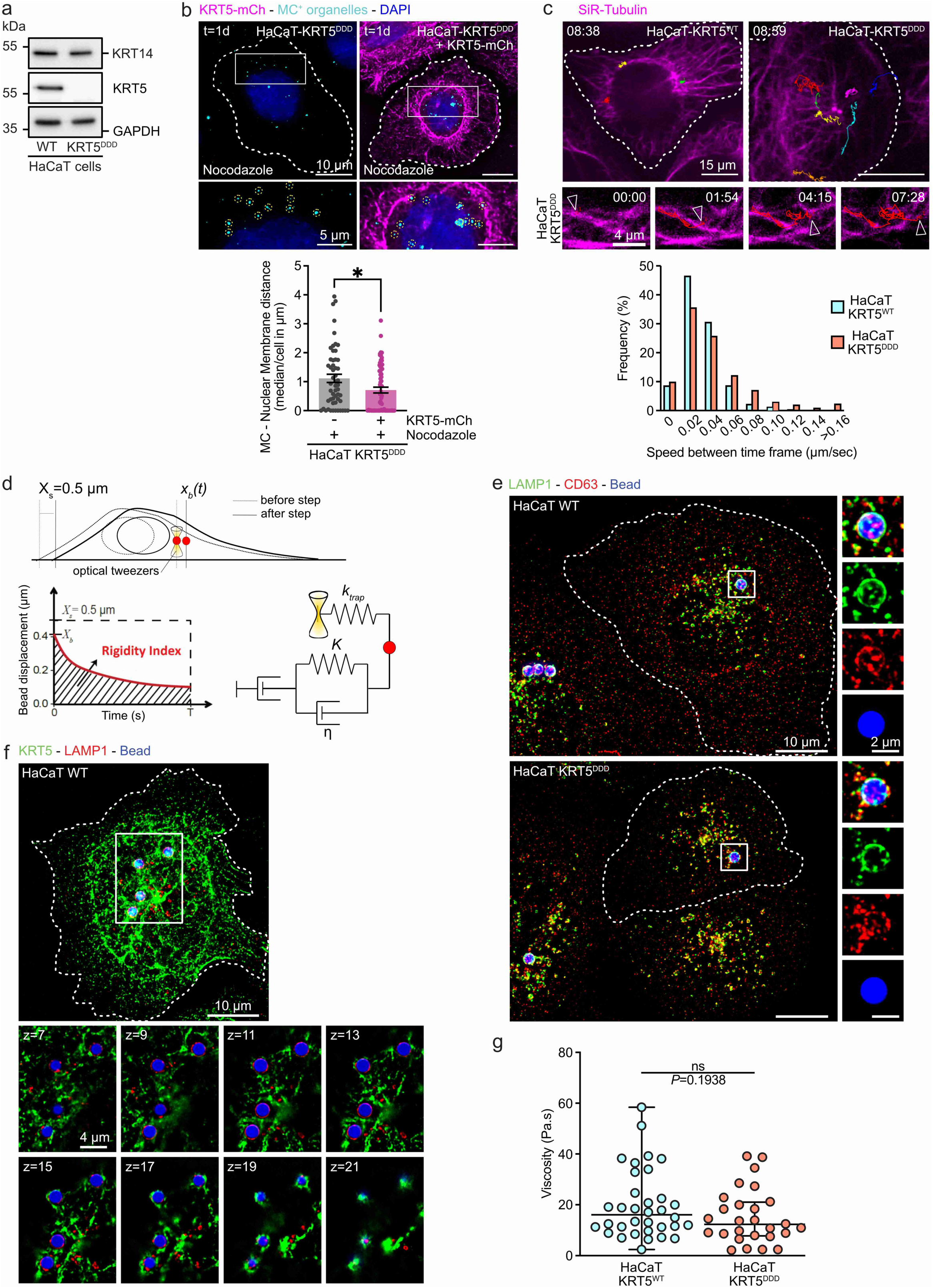
KRT5 expression is required for MC^+^ organelles perinuclear clustering and mechanical confinement. (**a**) Western blot of lysates of WT and KRT5^DDD^ HaCaT cells probed with anti-KRT14 (top), - KRT5 (middle) or -GAPDH (loading control, bottom) antibodies. (**b**) (Top) FM of nocodazole-treated KRT5^DDD^ HaCaT cells expressing (right) or not (left) KRT5-mCherry (magenta) 1 day after MCs deposition and stained with DAPI (blue). MCs captured 1 day after deposition by brightfield microscopy were pseudo-colored in cyan Automatically detected MCs are circled in yellow and cell outlines are delineated by dashed lines. (Bottom) Quantification of the median distance of MC^+^ organelles to the nucleus edge expressed per cell. (**c**) (Top) Live imaging frame of WT-(left) or KRT5^DDD^- (right) HaCaT cells incubated with SiR-Tubulin probe (magenta). MC^+^ organelles (not shown) were captured by brightfield, and their movements tracked throughout >8min of acquisition. The trajectories (colored lines) of detected MC^+^ organelles are shown. Arrowheads point the trajectory of a MC^+^ organelle over 7min in HaCaT-KRT5^DDD^ cells and its alignment along SiR-Tubulin^+^ MTs. (Bottom) Instantaneous speed in between time frames of tracked MC^+^ organelles in WT (cyan) or KRT5^DDD^ (red) HaCaT cells and presented as a frequency plot. (**d**) Diagram illustrating the microrheology experiment. (Top) A 2-µm-diameter bead internalized in the cell is trapped with an optical tweezer. At time *t* = 0 s, the microscope stage is moved in a *X_s_* = 0.5-µm step displacement. After the initial rapid displacement of the bead from the trap center, the bead position *x_b_*(*t*) relaxes towards the center of the optical trap, which acts as a spring. (Bottom left) Single particle tracking of the bead allows determination of the viscoelastic relaxation curves (see Fig. 5b). (Bottom right) Diagram of the Standard Linear Liquid (SLL) viscoelastic model. (**e**) IFM of WT (left) and KRT5^DDD^ (right) HaCaT cells having internalized beads (blue) and stained with anti-LAMP1 (green) and -CD63 (red) antibodies. (**f**) IFM of WT HaCaT cell having internalized beads (blue) and stained with anti-KRT5 (green) and -LAMP1 (red) antibodies. Insets show several z optical sections of the boxed area. (**f**) Quantification of the relaxation curves in Fig. 5b using the Standard Linear Liquid (SLL) viscoelastic model and analysis of the viscosity (in Pa.s) of the cytosolic microenvironment of the bead in WT (blue) or KRT5^DDD^ (red) HaCaT cells. (b, c, e, f) Cell outlines are delineated by dashed lines. Data are the average of at least three independent experiments presented as the mean ± SD or SEM. Two-tailed unpaired t test (b) and Mann-Whitney two-tailed unpaired t test (g) (ns, non-significant; *, *P* < 0.05).

**Extended Data Figure 6.**
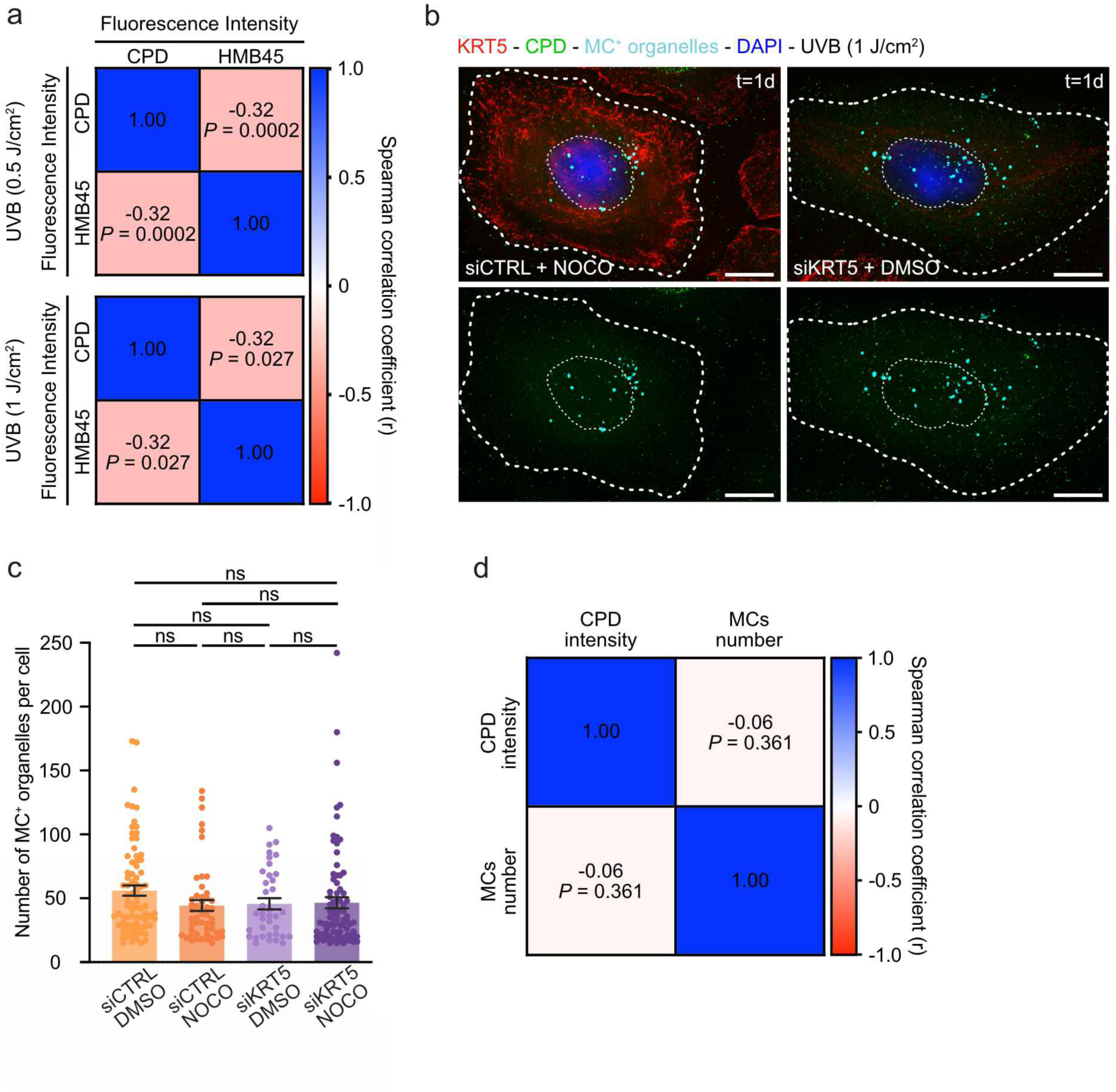
Perinuclear melanocore organelles have photoprotective activity. (**a**) Matrixes of correlation between the mean fluorescent nuclear intensities of CPD and HMB45 per HEK exposed to UVB doses of 0.5 J/cm^2^ (top) or 1 J/cm^2^ (bottom) analyzed in Fig. 6a-b and showing the non-parametric Spearman correlation coefficient (r). (**b**) IFM 1 day after MCs uptake of siCTRL-NOCO (left) or siKRT5-DMSO (right) treated HEKs exposed to 1 J/cm^2^ UVB dose before staining with anti-CPD antibody (green) and DAPI (cyan). MCs captured by brightfield microscopy were pseudo-colored in magenta. Cells are delineated by dashed lines. (**c**) Quantification of the mean number of MCs per cell treated in Fig. 6d. (**d**) Matrix of correlation between the mean nuclear CPD fluorescence intensity and the mean number of MCs per HEK analyzed in Figures 6 c-e and showing the non-parametric Spearman correlation coefficient (r). Data are the average of at least three independent experiments presented as the mean ± SEM. One-way ANOVA statistical analysis with Tukey post-hoc (ns, non-significant; ****, *P* < 0.0001).

